# Molecular determinants of selective and high-affinity binding of the scaffold protein PDZK1 to the transporter URAT1

**DOI:** 10.1101/2025.07.21.665923

**Authors:** E.V. Mymrikov, C. Wirth, J.I. Heinicke, J. Goll, B.A. Kern, C. Steck, A.K. Iaroslavtceva, T. Mühlethaler, A. Köttgen, C. Hunte

## Abstract

The renal solute carrier URAT1 (SLC22A12) is essential for urate homeostasis, with loss-of-function linked to renal hypouricemia, nephrolithiasis and lower gout risk. URAT1 function depends on binding the multi-PDZ domain scaffold protein PDZK1 (NHERF3), with a similar role suggested for the related NHERF1. The molecular basis of these interactions remains poorly understood. Using fluorescence anisotropy, we show that full-length human PDZK1 binds the C-terminal peptide of URAT1 with high affinity (*K*_D_ 170 nM), unlike NHERF1 (*K*_D_ >70 µM). The PDZ1 domain of PDZK1 alone is sufficient for high-affinity binding (*K*_D_ 160 nM), while PDZ4 provides a secondary site (*K*_D_ 1.35 µM), with both interactions characterized by rapid kinetics. Gel filtration shows that PDZK1 can bind two URAT1 peptides. X-ray structures of individual PDZ domains from PDZK1 and NHERF1 complexed with the URAT1 peptide reveal the molecular determinants for PDZK1’s higher affinity and selectivity. Murine Pdzk1 and Nherf1 bind Urat1 with high affinity indicating species-specific interactions. These data provide insights into URAT1 regulation by PDZ scaffold proteins with relevance for understanding urate homeostasis regulation and related disorders.

## INTRODUCTION

Solute carriers (SLCs) play a central role in excretion and reabsorption of metabolites in the kidney especially in the proximal tubule (Lewis et al., 2021, Lin et al., 2015, Zhang et al., 2019). It is increasingly recognized that SLCs require distinct interaction partners for full functionality (Frommelt et al., 2025), including scaffold proteins that contain multiple PSD-95/Dlg/ZO-1 (PDZ) domains, which are involved in regulation of their specific membrane localization (Biber et al., 2005, Brone & Eggermont, 2005, Walsh et al., 2015). Among those is the interaction between one of the major urate transporters, URAT1 (SLC22A12) and PDZK1 (NHE3 regulatory factor 3, NHERF3) (Leask et al., 2024, Nigam, 2018, Walsh et al., 2015, Wright et al., 2010). Common genetic variants in the *PDZK1* gene are associated with altered serum urate level (Ketharnathan et al., 2018, Kolz et al., 2009, Köttgen et al., 2013, Tin et al., 2018) and with gout susceptibility (Higashino et al., 2016, Major et al., 2024, Phipps-Green et al., 2016). Specifically, the genetic variant *rs1967017* in the *PDZK1* enhancer affects PDZK1 expression levels in the colon and small intestine and in model cell lines (Ketharnathan et al., 2018). *PDZK1* expression has been suggested to affect URAT1 localization at the brush border membrane of the proximal tubule cells (Ketharnathan et al., 2018, Leask et al., 2024, Nigam, 2018). Co-overexpression of PDZK1 and URAT1 in HEK293 cells resulted in the upregulation of urate uptake by transfected HEK293 cells (Anzai et al., 2004).

PDZ domains typically bind their target proteins via short, C-terminal linear and intrinsically disordered peptides, which are called PDZ binding motifs (PBM) (Amacher et al., 2020, Ivarsson, 2012). PDZK1 was initially identified as interactor of human URAT1 through a yeast two-hybrid screen using the C-terminal tail of URAT1 as a bait, and the interaction was confirmed by co-immunoprecipitation from co-overexpression in HEK293 cells as well as from membrane preparations of human kidney tissue (Anzai et al., 2004). Binding was attributed to three out of the four PDZ domains of PDZK1 (PDZ1, PDZ2, PDZ4) based on studies using fusion proteins containing the C-terminal tail of URAT1 (Anzai et al., 2004). Neither the binding affinity of full-length PDZK1 for URAT1 nor the precise contribution of the individual domains to this interaction are known.

NHERF1, the other PDZ domain scaffold protein abundant in human proximal tubule cells, was suggested to interact with URAT1, as binding of murine Nherf1 to Urat1 was demonstrated using co-immunoprecipitation (Cunningham et al., 2007) and blot overlay assays (Gisler et al., 2003). Yet, experimental data on the interaction of human URAT1 and NHERF1 are lacking. One should note that the C-terminal PBMs of the human and mouse URAT1 transporter differ (residues KSTQF and MSTRL, respectively), so that it would be interesting to know whether the interaction is species-specific.

In general, NHERF proteins interact via their PDZ domains with class I PBMs. The latter share the consensus sequence X^-4^-X^-3^-S/T^-2^-X^-1^-Φ^0^, where X is any amino acid at positions −1, −3 and −4, S/T^-2^ stands for serine or threonine at positions −2, and Φ^0^ is a hydrophobic residue at the very C-terminus (position 0) (Amacher et al., 2020, Lee & Zheng, 2010). The interaction of PDZ domains and PBM peptides has been intensively structurally characterized by X-ray and NMR analysis, with more than 40 structures of different human PDZ domains in complex with their respective target PBM peptides available (Amacher et al., 2020). Despite multiple, functionally relevant interactions described between PDZ domain proteins and human SLCs (Biber et al., 2005, Frommelt et al., 2025, Gisler et al., 2003, Walsh et al., 2015), only a single structure of a PDZ domain in complex with the PBM peptide of a human SLC is available, namely of the PDZ4 domain of PDZK1 with SLC15A2 (PEPT2 transporter) (Hajizadeh et al., 2018). Thus, structural information regarding how PDZ domains recognize and bind PBMs of SLCs, including URAT1, is scarce. Understanding the molecular basis of these interactions is important, especially for multi-domain scaffold proteins such as PDZK1 and NHERF1. Structural data are needed to identify the specific determinants that govern domain selectivity and binding affinity, which likely correlate with distinct regulatory functions.

Insights into PDZ domain/PBM specificity have been gained by phage-display peptide library screening for individual PDZ domains, which enabled the classification of PDZ domains into distinct subclasses and identification of their preferred PBMs (Ivarsson et al., 2014, Teyra et al., 2020, Tonikian et al., 2008). Comparative structural analysis of representative PDZ domains with different specificity in complex with respective PBM peptides revealed key amino acid positions within the PDZ domains that were linked to specificity for defined PBMs (Ernst et al., 2014). A systems biology approach using high-throughput holdup assays with maltose-binding protein-fused PDZ domains has enabled the quantification of binding affinity and specificity across a substantial part of the human PDZome, including NHERF1 and PDZK1 domains, covering a broad range of affinities from sub-micromolar to hundreds of micromolar (Gogl et al., 2022). Despite this progress, the molecular basis that underlies the distinct selectivity of a PBM for a specific PDZ domain(s), especially in the case of multi-domain scaffold proteins such as NHERF1 and PDZK1, as well as the structural determinants controlling the affinity of the individual interaction remain poorly understood. Such knowledge would be also important to understand the higher order organization of SLCs to supramolecular associations in the membrane. An example for such association is the complex of the sodium/glucose cotransporter SGLT2 (SLC5A2) with PDZK1IP1 (MAP17) (Cui et al., 2023), which is essential for transport activity (Coady et al., 2017).

As a multi-domain protein, PDZK1 bears potential to associate different proteins, as it not only interacts with URAT1 but with more than ten other renal transporters including the name-giving sodium-hydrogen exchanger 3 (NHE3, SLC9A3) and the sodium-dependent phosphate transporter 2A (NaPi-IIa, SLC34A1) as shown by yeast two-hybrid screening, pull-down and/or co-immunoprecipitation assays (Cha et al., 2017, Gisler et al., 2003, Kato et al., 2005, Miyazaki et al., 2005, Rossmann et al., 2005, Srivastava et al., 2019, Thomson et al., 2005). PDZK1 has been proposed to simultaneously bind different transporters, forming so-called “urate transportosomes” (Anzai et al., 2007, Prestin et al., 2017, Wright et al., 2010) or “urate transportasomes” (Dalbeth & Merriman, 2009, Leask et al., 2024, Reginato et al., 2012), which could optimize urate transport across the membrane.

In this study, we provide a detailed characterization of the regulatory interaction between the medically relevant renal transport protein URAT1 with the multi-PDZ domain scaffold proteins PDZK1 and NHERF1. With biochemical and biophysical methods using purified full-length proteins and their individual domains, we show that human PDZK1 binds URAT1 with high-affinity and rapid kinetics via PDZ1 and PDZ4 domains, whereas the other two domains do not contribute substantially, and that the interaction with NHERF1 is very weak. The two high-affinity binding sites of PDZK1 could enable the formation of higher-order complexes, supported by results proving the simultaneous binding of two URAT1 peptides to full-length PDZK1. We reveal the underlying structural basis with high-resolution X-ray structures of the PDZ domain/PBM complexes, showing that the number of direct side chain interactions correlates with affinity. We also demonstrate that the interaction between URAT1 and the scaffold proteins is species-specific, as for both murine Pdzk1 and Nherf1, high affinity binding of Urat1 peptide was observed. These data elucidate the molecular and structural basis of URAT1 regulation by PDZ scaffold proteins important to understand the mechanism of urate transport and its dysfunction, and sheds light on the organization SLC assemblies at the membrane.

## RESULTS

### Human full-length PDZK1 but not NHERF1 strongly binds to the C-terminus of URAT1

In order to gain precise quantitative information about the functionally relevant interaction of PDZK1 with the human renal solute carrier URAT1 (Anzai et al., 2004, Ketharnathan et al., 2018, Nigam, 2018, Tin et al., 2018) and to challenge the postulated interaction with NHERF1, we designed constructs of full-length PDZK1 and NHERF1, as well as of their individual PDZ domains, for recombinant protein production (Fig. 1A). We purified independently three biological replicates of each construct (Fig. 1B and Fig. S1) and used a fluorescent anisotropy (FA) assay for *K*_D_ determination with fluorescently labelled peptide that correspond to the 15 C-terminal residues of URAT1 (‘URAT1 peptide’). As the binding of NaPi-IIa to NHERF1 has been characterized in detail (Levi et al., 2019, Mamonova et al., 2015) and this transporter has been shown to interact with mouse and rat PDZK1 (Gisler et al., 2003, Gisler et al., 2001, Hernando et al., 2002), we included the peptide that corresponds to the 15 C-terminal residues of NaPi-IIa (‘NaPi-IIa peptide’) for comparison. We observed an increase of FA signal with an increase in the PDZK1 concentration in the assay, demonstrating that full-length PDZK1 binds both, the URAT1 and NaPi-IIa peptides (Fig. 2A). Notably, PDZK1 showed a distinct preference for URAT1, binding the respective peptide with *K*_D_^app^ of 0.17 ± 0.01 µM. Its affinity is much lower for the NaPi-IIa peptide, with almost 50 times higher *K*_D_^app^ (8.1 ± 0.5 µM). We refer to *K*_D_^app^ for full-length PDZK1 and NHERF1 as they have four and two PDZ domains, respectively, each of them is a potential PBM binding site. In contrast to PDZK1, NHERF1 binds the NaPi-IIa peptide with *K*_D_^app^ of 4.0 ± 0.2 µM, much stronger than the URAT1 peptide (Fig. 2B). This value is very similar to the affinity of PDZK1 for the NaPi-IIa peptide. NHERF1 binding to the URAT1 peptide is very weak, and the saturation was not reached in this case. The estimated *K*_D_^app^ value for the NHERF1/URAT1 interaction was above 70 µM. Taken together, URAT1 strongly binds to PDZK1 but weakly to NHERF1. We therefore suggest PDZK1 as physiological binding partner of URAT1 in the human kidney and oppose the interaction of URAT1 with NHERF1, which was previously deduced from mouse samples (Cunningham et al., 2007, Gisler et al., 2003).

**Figure 1.**
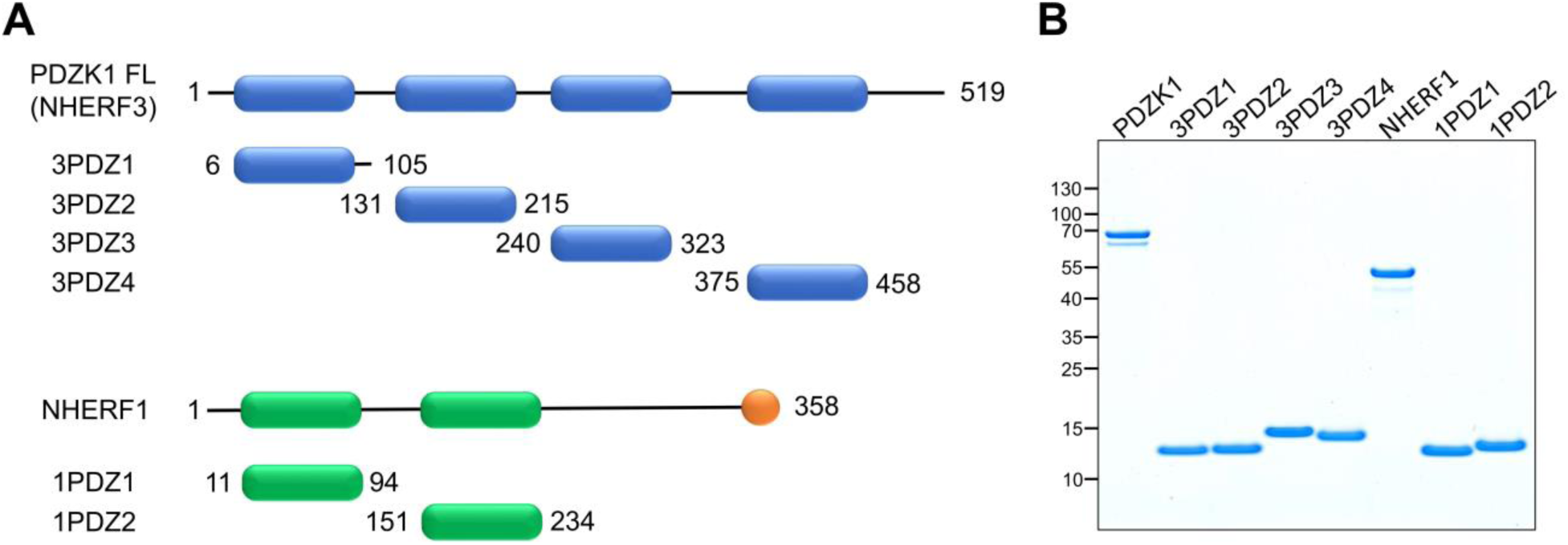
Full-length PDZK1 and NHERF1 as well as individual domains used. (**A**) Schematic representation of full-length proteins and individual PDZ domains (blue and green colored boxes). The start and end of each construct are indicated by respective amino acid residue number. PDZ domains of PDZK1 are referred to as 3PDZ1 to 3PDZ4, those of NHERF1 as 1PDZ1 and 1PDZ2. NHERF1 carries a C-terminal ezrin-binding domain depicted as an orange circle. (**B**) Quality control of purified full-length proteins and PDZ domains by SDS-PAGE analysis. Representative Coomassie-stained gel with positions of co-separated molecular mass standard proteins (kDa) indicated on the left. Analysis of biological replicates is shown in Fig. S1.

**Figure 2.**
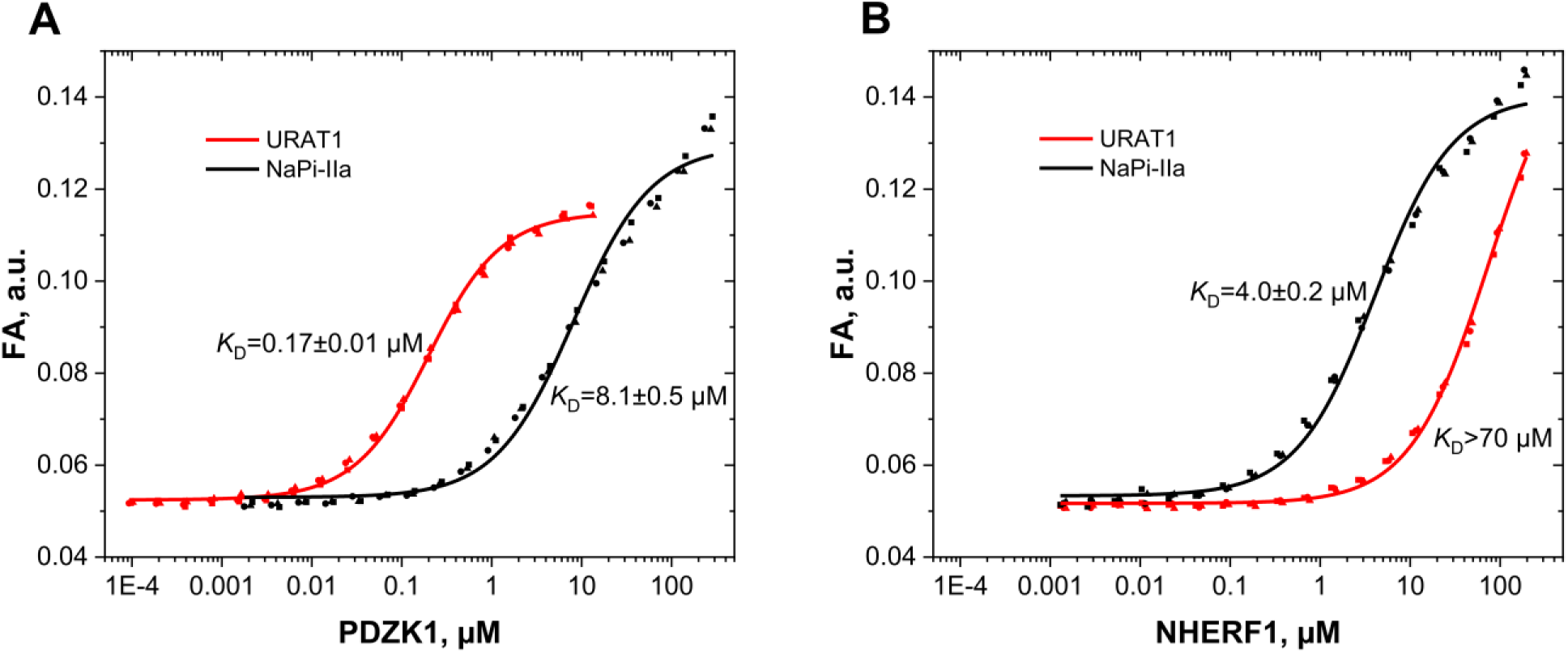
Human full-length PDZK1, but not NHERF1 binds the URAT1 peptide with a nanomolar affinity. Binding affinities (*K*_D_^app^) of full-length PDZK1 (**A**) and NHERF1 (**B**) for the fluorescently labeled C-terminal URAT1 (black) and NaPi-IIa (red) peptides were determined with the fluorescence anisotropy assay. Each measurement was done with three independent biological replicates (data points of different shape) and a concatenated fit was performed (solid lines) for *K*_D_^app^ determination. Respective *K*_D_^app^ ± SE values indicated as “*K*_D_” are shown.

### Individual PDZ domains of PDZK1 and NHERF for the C-terminus of URAT1 have distinct affinities ranging from high nanomolar to high micromolar

In order to identify which domain is predominantly responsible for the interaction with the above-mentioned transporters, we determined the affinities of each individual PDZ domain for URAT1 and NaPi-IIa peptides using the FA assay and three biological replicates (Fig. 3). Hereafter, PDZ domains of PDZK1 (NHERF3) are referred to as 3PDZ1, 3PDZ2, 3PDZ3, and 3PDZ4, and those of NHERF1 as 1PDZ1 and 1PDZ2 (Fig. 1A). We produced and purified all six individual PDZ domains of PDZK1 and NHERF1, each of which had a C-terminal His_8_-tag except for 3PDZ1, for which the N-terminal His_8_-tag was removed (Fig. 1, Fig. S1 C-E). In all measurements, an increase of fluorescence anisotropy was observed, indicating at least a weak binding. For *K*_D_ values below 25 µM, reflecting high to medium affinity binding, the saturation plateau was reached and *K*_D_ values were determined precisely (Table 1). In cases of weak binding affinity, the saturation plateau could not be reached (see e.g. Fig. 3C), as protein concentrations above about 500 µM non-specifically affect the fluorescence anisotropy readout. For such low-affinity interactions, an approximation of *K*_D_ values is provided.

**Figure 3.**
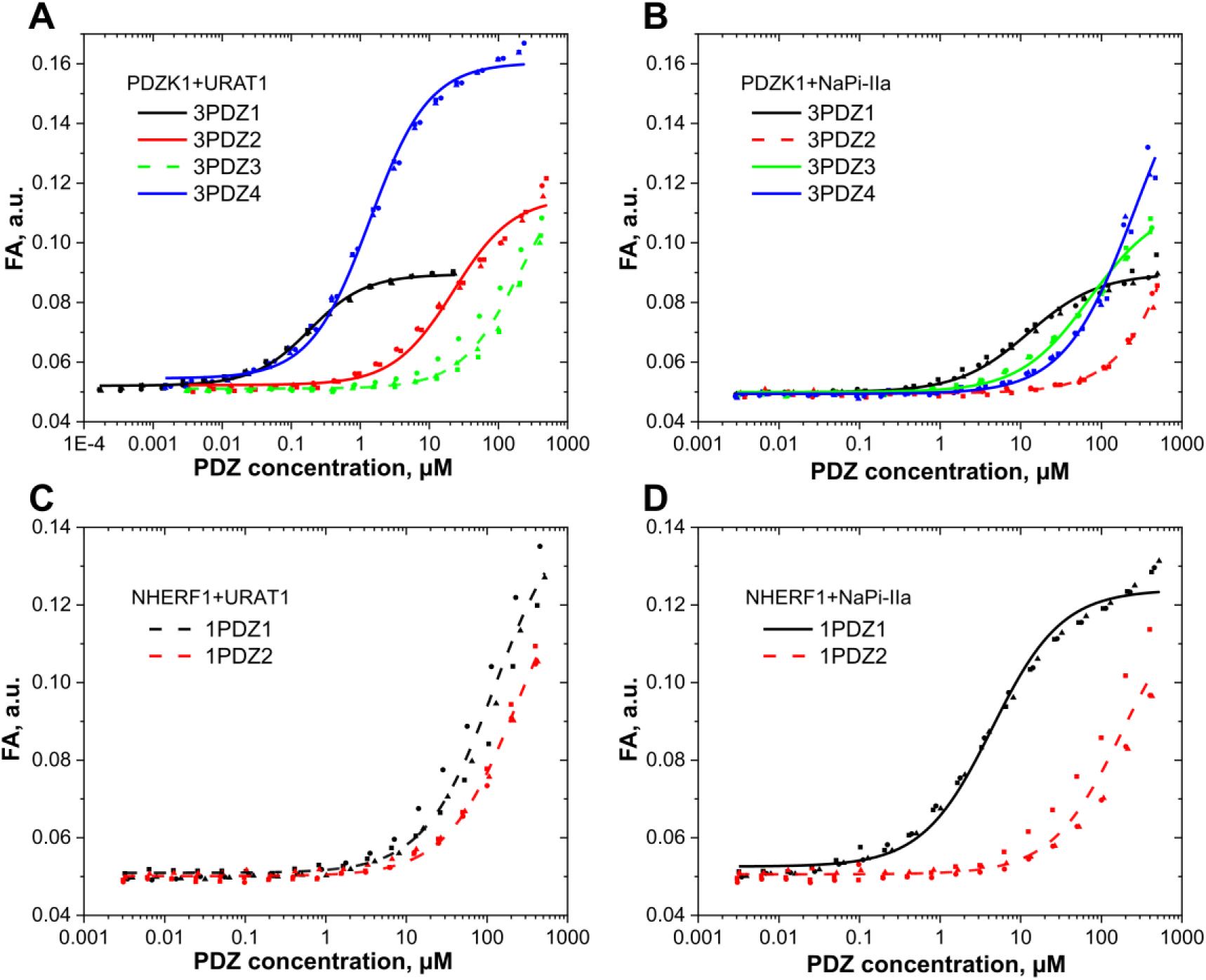
Binding affinities (*K*_D_) of individual PDZ domains of human PDZK1 and NHERF1 for URAT1 and NaPi-IIa peptides determined with FA assay. (**A**) PDZ domains of PDZK1 with URAT1 peptide; (**B**) PDZ domains of PDZK1 with NaPi-IIa peptide; (**C**) PDZ domains of NHERF1 with URAT1 peptide; (**D**) PDZ domains of NHERF1 with NaPi-IIa peptide. Each measurement was performed with three independent biological replicates (data points of different shape) and a concatenated fit was performed for *K*_D_ determination. If the saturation plateau has not been reached and we could only estimate *K*_D_ values, the fitting curve is shown as a dashed line. All *K*_D_ values are summarized in Table 1.

**Table 1.** Binding affinities (*K*_D_ in µM) of PDZ domains from PDZK1 and NHERF1 for URAT1 and NaPi-IIa peptides (PBMs are in brackets) determined with FA assay. *K*_D_^app^ values of full-length proteins are shown for comparison. Values are *K*_D_ ± SE of the fit.

|  |  | URAT1 (KSTQF) | NaPi-IIa (NATRL) |
| --- | --- | --- | --- |
| PDZK1 | full-length | $0.17 \pm 0.01$ | $8.1 \pm 0.5$ |
| | 3PDZ1 | $0.16 \pm 0.01$ | $12.5 \pm 0.9$ |
| | 3PDZ2 | $22.1 \pm 1.9$ | n.d.** |
| | 3PDZ3 | $>200^*$ | $>70^*$ |
| | 3PDZ4 | $1.35 \pm 0.06$ | $>260^*$ |
| NHERF1 | full-length | $>70^*$ | $4.0 \pm 0.2$ |
| | 1PDZ1 | $>120^*$ | $4.4 \pm 0.3$ |
| | 1PDZ2 | $>230^*$ | $>170^*$ |

\*estimated $K_D$ values as saturation was not reached
\*\*too low affinity to approximate $K_D$ value

3PDZ1 domain binds both URAT1 and NaPi-IIa peptides with the highest affinity among all four domains (0.16 ± 0.01 µM and 12.5 ± 0.9 µM, respectively) (Fig. 3A, B and Table 1). Affinities of 3PDZ1 for URAT1 and NaPi-IIa peptides are very close to those of full-length PDZK1 (Table 1), indicating that this domain is sufficient for the high affinity interaction with both peptides and allosteric effects can be excluded. In addition, 3PDZ4 also binds strongly the URAT1 peptide with *K*_D_ of 1.35 ± 0.06 µM, which is 8.5-fold higher the *K*_D_ value of 3PDZ1. The URAT1 affinity for 3PDZ2 is lower with a *K*_D_ of 22.1 ± 1.9 µM (140-fold higher as compared to 3PDZ1). This domain hardly interacts with the NaPi-IIa peptide, while 3PDZ4 binds it very weakly (Fig. 3B and Table 1). 3PDZ3 binds both peptides with very low affinity and can be excluded from contributing to interactions with URAT1 and NaPi-IIa. In line with the very weak interaction of full-length NHERF1 and the URAT1 peptide shown above, the individual PDZ domains of both proteins also demonstrated only moderate binding to the respective peptide (Fig. 3C and Table 1). As a control, we determined the affinity for the interaction between 1PDZ1 and the NaPi-IIa peptide with a *K*_D_ of 4.40 ± 0.27 µM and showed very weak binding of 1PDZ2 to this peptide (*K*_D_>170µM), in good agreement with previously published data (Mamonova et al., 2015) (Fig. 3D and Table 1). Taken together, the data suggest that PDZK1 can bind both URAT1 and NaPi-IIa through its PDZ1 domain with a higher affinity for URAT1, whereas NHERF1 can bind NaPi-IIa but not URAT1. In addition, as the PDZ4 domain of PDZK1 also shows substantial affinity for the URAT1 peptide, full-length PDZK1 may have the capacity to bind two URAT1 molecules. As we have shown that the individual PDZ domains have distinct binding affinities for the URAT1 peptide ranging from high nanomolar to high micromolar *K*_D_ values, these domains provide an ideal example to dissect the underlying structural basis of such affinity and selectivity.

### The canonical features of URAT1 PBM peptide binding to individual PDZ domains

In order to precisely describe the structural basis underlying the broad range of affinities for URAT1 binding to the individual PDZ domains of PDZK1, we determined X-ray structures of three domains with the URAT1 PBM peptide (KSTQF) bound, covering high (3PDZ1 and 3PDZ4) to low (3PDZ2) binding affinity. We crystallized, in various conditions, each PDZ domain with the PBM peptide as C-terminal fusion to enable complex formation between protomers (Table S2).

For comparison, a similar approach was applied to PDZ1 domain of NHERF1 (1PDZ1), which binds the URAT1 peptide with very low affinity. For each construct, a high-resolution X-ray diffraction structure was obtained with resolutions between 1.2 – 1.45 Å (Table S3).

Thanks to the high resolution in four obtained structures (3PDZ1^KSTQF^, 3PDZ2^KSTQF^ 3PDZ4^KSTQF^, and 1PDZ1^KSTQF^), PDZ domains and bound PBM peptide as well as many solvent molecules could be modeled unambiguously. The refined structures comprise nearly all residues (Table S3). Noteworthy, in all structures, the PDZ domains bind the PBM peptide from a neighboring protomer. This resulted in variety of crystalline arrangements that are similar to those previously observed for other PDZ domain/PBM fusion constructs (Elkins et al., 2007). The PDZ domains feature the canonical globular PDZ domain fold (Amacher et al., 2020, Hung & Sheng, 2002, Skelton et al., 2003, Songyang et al., 1997) and comprise six β-strands and two α-helices with the secondary structure topology β1-β2-β3-α1-β4-β5-α2-β6 (Fig. 4A, D). The 3PDZ1 domain has an additional helix α3 followed by a short loop that do not change the core domain architecture (Fig. 4E). This extension of 15-residues was required to obtain a stable recombinant protein.

**Figure 4:**
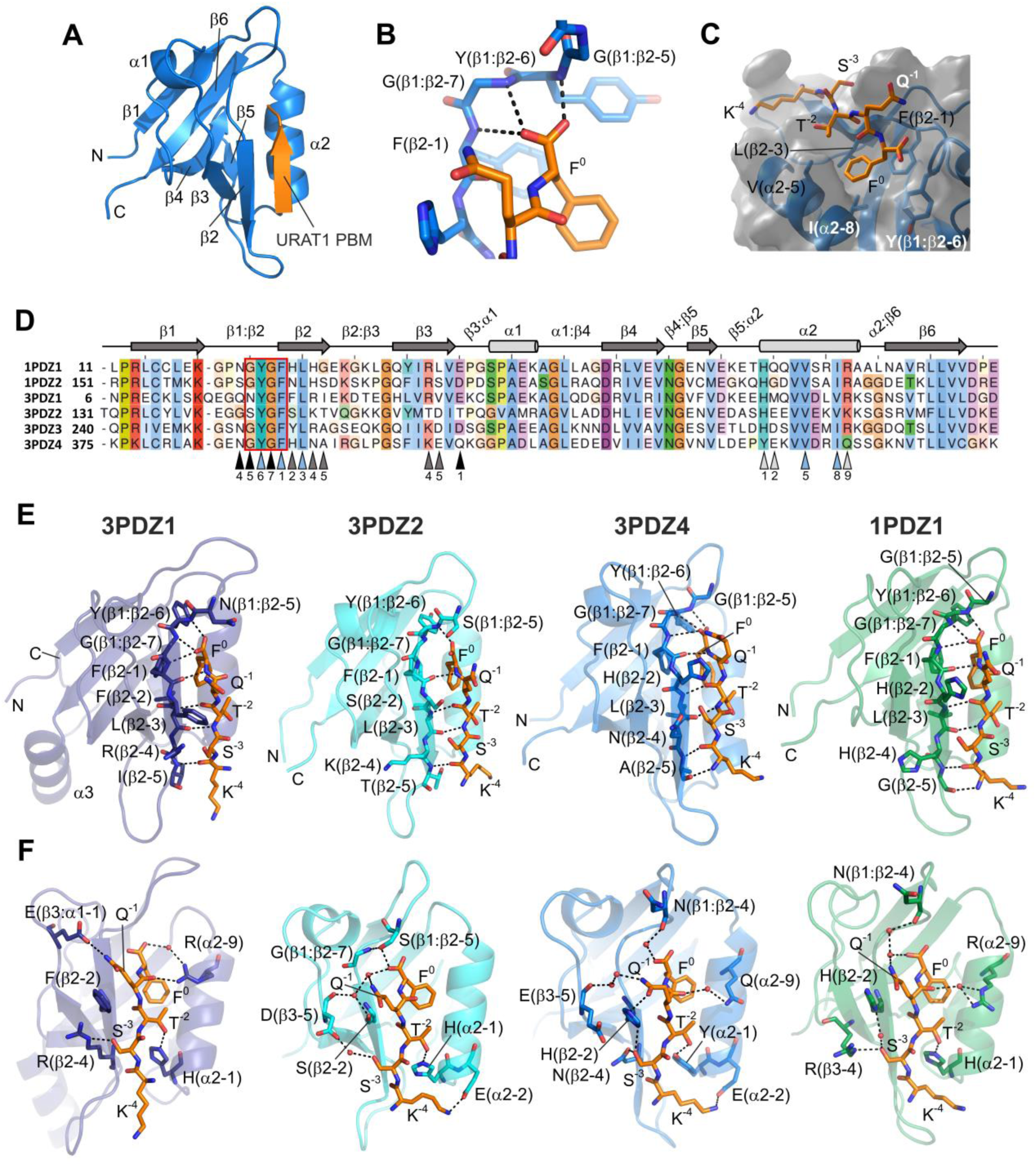
X-ray structures of PDZK1 and NHERF1 PDZ domains with bound URAT1 PBM peptide (KSTQF). (**A**) Overall structure of the PDZ^KSTQF^ (3PDZ4^KSTQF^ as an example) in cartoon representation. The PDZ domain is in blue and the PBM peptide forming a β-strand is orange. Secondary structure elements as well as N- and C-termini of the PDZ domain are indicated. (**B**) The carboxylate binding loop of 3PDZ4 domain that embraces the C-terminal carboxylate of the PBM peptide. Residues of the conserved XYGF motif and F^0^ of the PBM are shown as sticks. (**C**) The hydrophobic pocket of 3PDZ4 domain that accommodates the F^0^ side chain of the PBM peptide. The conserved residues lining the surface of this pocket are indicated, whereas main chain atoms of the PDZ domain are omitted. (**D**) Multiple sequence alignment of human NHERF1 and PDZK1 PDZ domains with indication of secondary structure elements and positions of residues that contribute to the binding of the URAT1 PBM peptide. Conserved secondary structure elements are schematically depicted on top; positions of the residues (naming according to (Skelton et al., 2003)) involved in the interaction with the PBM peptide are marked below the alignment. Black triangles point to residues in loops, dark grey triangles indicate residues in β-strands, light grey triangles indicate residues in α-helices, and blue triangles show hydrophobic residues lining the surface of the hydrophobic pocket (see **C**). Conserved XYGF motif coordinating C-terminal carboxylate is outlined with a red rectangle. Sequence alignment was done in Jalview (2.11.3.3 version) (Waterhouse et al., 2009) using ClustalO algorithm. (**E, F**) Cartoon representation of 3PDZ1^KSTQF^ (deep blue), 3PDZ2^KSTQF^ (cyan), 3PDZ4^KSTQF^ (blue), and 1PDZ1^KSTQF^ (green) structures, the URAT1 PBM peptide is shown as orange sticks. (**E**) PDZ domain residues involved in the hydrogen bond network (black dotted lines) between main chain atoms of the PDZ domain and the PBM peptide are shown as sticks. These interactions contribute to the coordination of C-terminal carboxylate and β-sheet extension. (**F**) Specific interactions involving the side chains of PDZ domain residues (shown as sticks) are highlighted as black dotted lines. These interactions are likely to be determinants for the specific binding of the URAT1 PBM peptide. Water molecules involved in the coordination of the PBM residues are depicted as red spheres.

In line with the canonical binding mode (Amacher et al., 2020), the PBM peptide binds as anti-parallel β-strand in a cleft formed by the β2-strand and the α2-helix extending the β2-β3 sheet of the PDZ domains (Fig. 4A, E). A typical antiparallel β-strand H-bond network is present and very similar in all four structures, except for the main chain nitrogen atom of K^-4^, which is the first atom of the PBM peptide (Fig. 4E, Fig. 5).

**Figure 5.**
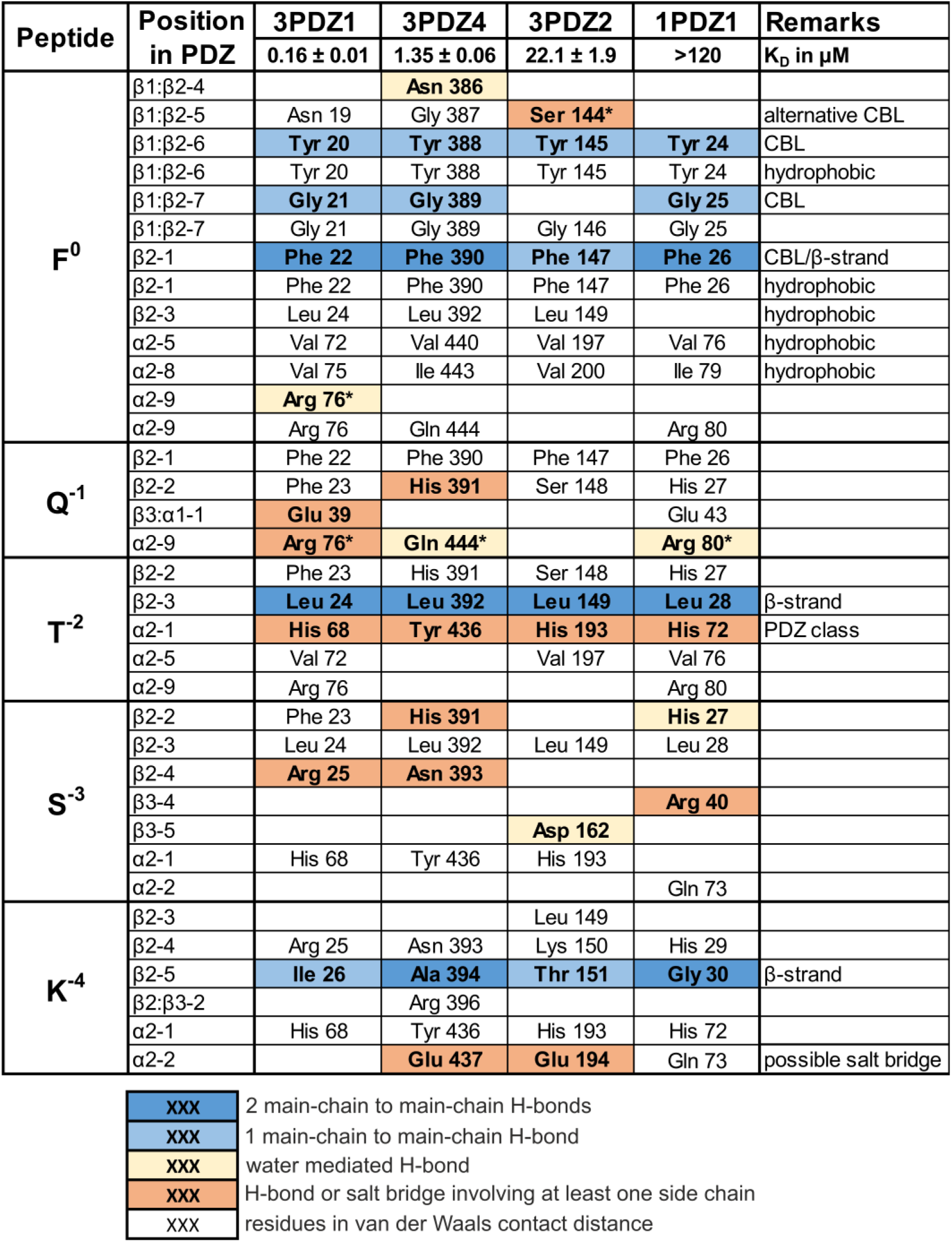
Interaction analysis highlights importance of direct H-bonds for high-affinity binding between URAT1 peptide and PDZ domains of PDZK1 and NHERF1. All PDZ residues in a distance shorter than 4 Å from the respective PBM residue (the first column) as well as those coordinating the PBM peptide via water-mediated H-bonds are shown. Residues in bold are involved in H-bonds or salt bridges, while residues in van der Waals contact distance are in regular font. The cell is blank if the PDZ residue at the corresponding position does not interact with the PBM residues. The “Remarks” column highlights interactions of the carboxylate binding loop (“CBL”), in the hydrophobic pocket (“hydrophobic”) and of the β-sheet extension (β-strand). Residue at α2-1 position coordinates the T^-2^ and was suggested to be a hallmark for a PDZ class. Asterisk (*) indicates contacts between a side chain of a PDZ domain residue and a main chain atom of the PBM residue, which therefore are not specific to a PBM sequence.

The second typical feature is the carboxylate binding loop (β1:β2) of the PDZ domains with its conserved XYGF motif (Fig. 4B, D). It encompasses and coordinates the C-terminal carboxylate of the PBM peptide. In the following, the residues of the PDZ domains are named according to their position in the canonical secondary structure (see Fig. 4D) as introduced by Skelton and colleagues (Skelton et al., 2003). In the structures of the complexes 3PDZ1^KSTQF^, 3PDZ4^KSTQF^, and 1PDZ1^KSTQF^, the coordination of the C-terminal carboxylate is realized through three polarized H-bonds contributed by backbone N-atoms of residues Y(β1:β2-6), G(β1:β2-7), and F(β2-1) of the XYGF motif (Fig. 4B, E). This type of carboxylate coordination has also been called nest carboxylate binding structural motif (Watson & Milner-White, 2002). In contrast, the coordination of the carboxylate in the structure 3PDZ2^KSTQF^ has only one direct H-bond to the main chain of the XYGF motif (Fig. 4E). The carboxylate is slightly rotated as compared to its orientation in the other structures of this study forming a single H-bond with the side chain of S(β1:β2-5), which bridges the carboxylate to G(β1:β2-7) (Fig. 4F). This partial and unusual coordination of the carboxylate could contribute to the lower affinity of 3PDZ2 for the URAT1 peptide.

The third component is the binding of the side chain of the terminal residue (Φ^0^) in the hydrophobic pocket formed by residues of the β2-strand and the α2-helix. So far, known structures of PDZ domains of NHERF proteins in complex with PBM peptides were limited to those with leucine as Φ^0^, with one exception of 3PDZ2 in a complex with the artificial peptide IKITKF (Ernst et al., 2014). In our four structures with the natural URAT1 PBM peptide, the side chain of F^0^ is surrounded by four hydrophobic residues Y(β1:β2-6), F(β2-1), L(β2-3), V(α2-5) that are strictly conserved in PDZK1 and NHERF1 PDZ domains (Fig. 4C). I(α2-8) in 3PDZ4 and 1PDZ1 or V(α2-8) in 3PDZ1 and 3PDZ2 completes the hydrophobic pocket (Fig. 4C). Notably, when the shorter side chain of valine is present in α2-8 position, the L(β2-3) side chain is rotated towards the F^0^ side chain. In contrast, the I(α2-8) in 3PDZ4 and 1PDZ1 seems to force the side chain of L(β2-3) to rotate away from the F^0^ side chain. These variations, however, are most likely not decisive for the observed differences in affinities, as both I(α2-8) and V(α2-8) are present in PDZ domains that bind the URAT1 peptide with high and low affinities. Multiple conserved non-polar interactions contribute to the stabilization of the PBM peptide in the hydrophobic pocket (Fig. 5). Another characteristic feature of the PDZ domain/PBM binding mode is the stabilization of the side chain of the residue in −2 position of the PBM. In all structures determined in this study, the side chain of T^-2^ of the PBM peptide is pointing toward the binding cleft formed by the β2-strand and the α2-helix and interacts with the side chain of the residue in position α2-1 (Fig. 4F). In the 3PDZ1, 3PDZ2, and 1PDZ1 domains, a histidine residue in the α2-1 position is providing an H-bond to the threonine side chain. In contrast, the H-bond partner is a tyrosine residue in that position in the 3PDZ4 domain (Fig. 4D).

Taken together, all PDZ^KSTQF^ structures determined in the present study demonstrate overall similar interaction patterns with little variations: for the β-sheet extension, the carboxylate binding loop, the binding mode of F^0^ to the hydrophobic pocket and the coordination of the T^-2^ side chain by the α2-1 residue (Fig. 5). These features are present in most known structures of class I PDZ domains, that bind PBMs with threonine at −2 position such as the URAT1 PBM. Therefore, the main determinants for the wide range of affinities for the individual PDZ domains must be contributed by domain-specific interactions.

### Specific interactions between PDZ domains and the URAT1 PBM peptide

In contrast to the canonical features described above, three other residues of URAT1 PBM (Q^-1^, S^-3^ and K^-4^) have a unique binding mode for each of the PDZ domains analyzed (Fig. 4F). This implies that mostly these interactions are decisive for selectivity and affinity for the individual domain. The side-chains of these three PBM residues are surface-exposed in contrast to aforementioned F^0^ and T^-2^, which both point towards the cleft between the β2-strand and the α2-helix. Multiple non-polar interactions to each residue of the PBM peptide contribute to the individual binding mode (Fig. 5). Further, the side chain of Q^-1^ is coordinated directly by the carboxylate of E(β3:α1-1) in the 3PDZ1^KSTQF^ structure (Fig. 4F), and F(β2-2) is in van der Waals contact distance with the side chain of Q^-1^ stabilizing its orientation for optimal H-bond formation with E(β3:α1-1). In the 3PDZ4^KSTQF^ structure, Q^-1^ forms a direct H-bond with the imidazole moiety of H(β2-2). In addition, it is connected to the main chain carbonyl oxygen of E(β3-5) via two water molecules (Fig. 4F). In the 1PDZ1^KSTQF^ structure, in which E(β3:α1-1) and H(β2-2) are also present, Q^-1^ interacts with none of these residues but is coordinated by N(β1:β2-4) via water molecules. Similar to the 1PDZ1^KSTQF^ structure, the coordination of Q^-1^ in the 3PDZ2^KSTQF^ structure lacks a direct H-bond, but instead is achieved via water molecules to D(β3-5) and S(β2-2) residues.

The side chain of S^-3^ is coordinated by the ζ-nitrogen atom of R(β2-4) in 3PDZ1^KSTQF^ structure. Interestingly, the aforementioned F(β2-2) forms a cation-π interaction with the guanidinium group of R(β2-4), thereby orienting also the side chain of the latter for stable binding of the PBM peptide. Interestingly, in 3PDZ4^KSTQF^ structure, the residues at the same positions are also coordinating the S^-3^ side chain. Both N(β2-4) and H(β2-2) form an H-bond with the γ-oxygen of S^-3^ (Fig. 4F). In 1PDZ1, H(β2-2) is also involved in S^-3^ coordination. However, in this case the H-bond is mediated by a water molecule. Notably, in the 1PDZ1^KSTQF^ structure, the R(β3-4) contributes a direct H-bond to S^-3^, structurally replacing the residue at the position β2-4 in 3PDZ1^KSTQF^ and 3PDZ4^KSTQF^ structures (Fig. 4F). In the 3PDZ2^KSTQF^ structure, the coordination of side chain of S^-3^ is less specific as this residue forms a single water-mediated H-bond to the side chain of D(β3-5).

The protonable K^-4^ forms an ion pair with the side chain of E(α2-2) in the 3PDZ2^KSTQF^ and 3PDZ4^KSTQF^ structures (Fig. 4F). This electrostatic interaction appears to be weaker in the 3PDZ2 ^KSTQF^ structure, as the side chain position of K^-4^ side is poorly resolved. The 3PDZ1 and 1PDZ1 domains have uncharged residues at the α2-2 position, which do not interact with K^-4^ in the respective structures (Fig. 4D, F). Interactions described above for Q^-1^, S^-3^ and K^-4^ residues involve their side chains and thus are specific for the URAT1 PBM peptide. In three of the structures, a further interaction between the peptide and the respective domain involves the residue at position α2-9 that forms H-bond(s) with main chain atoms of the PBM peptide (Fig. 4F). In the 3PDZ1^KSTQF^ structure, R(α2-9) forms a direct H-bond with the main chain of Q^-1^ and a water-mediated H-bond with the C-terminal carboxylate (Fig. 4F). Q(α2-9) in the 3PDZ4^KSTQF^ and R(α2-9) in the 1PDZ1^KSTQF^ structure, form a water-mediated H-bond with the main chain of Q^-1^ (Fig. 4F). In the 3PDZ2^KSTQF^ structure, lysine at α2-9 position does not bind to the PBM peptide (Fig. 4D, F). This interaction with the main chain of the PBM peptide is likely to contribute to and modulate the binding affinity.

Taken together, the four structures described above, which represent complexes with binding affinities in the range of high nanomolar to high micromolar, show specific coordination signatures with respect to the interaction between the Q^-1^, S^-3^ and K^-4^ side chains and the residues of the individual PDZ domains. Whereas the canonical binding features show little variations, side-chain-mediated polar interactions appear to be domain-specific, with more polar interactions between the URAT1 PBM peptide and 3PDZ1 and 3PDZ4, the higher affinity binders, as compared to the lower affinity binders 3PDZ2 and 1PDZ1. Complexes of 3PDZ1 and 3PDZ4 are characterized by more direct H-bonds mediated by side chains of the respective PDZ domain to the URAT1 PBM peptide and especially to residues Q^-1^ and S^-3^ (Fig. 5). In contrast, 3PDZ2 and 1PDZ1 domains, which bind the URAT1 peptide with lower affinity, present less of such H-bonds. Notably, in the two respective structures, several water molecules bridge the gaps between Q^-1^ and S^-3^ and side chains of the PDZ domain residues suggesting sub-optimal fit of the URAT1 PBM to the architecture of 3PDZ2 and 1PDZ1 binding sites.

### Contribution of the PBM residues at −5 and −6 position to the interaction with the PDZ domains

Next, we questioned whether URAT1 residues located further upstream from the C-terminus (V^-5^ and L^-6^) might contribute to the high-affinity binding of 3PDZ1 and 3PDZ4 as it was described for some PDZ domains (Tonikian et al., 2008). Thus, we fused the 7-mer URAT1 PBM peptide (VLKSTQF) to 3PDZ1 and 3PDZ4 domains and crystallized the respective constructs. The structures 3PDZ1^VLKSTQF^ and 3PDZ4^VLKSTQF^ were determined at slightly lower resolution (Table S4) compared to the 3PDZ1^KSTQF^ and 3PDZ4^KSTQF^ structures. In both structures, the binding modes of VLKSTQF are overall very similar as compared to those with the shorter PBM peptide (Fig. 4E, F and Fig. S2, S3). Further, there are only minor contributions of the V^-5^ and L^-6^ side chains to the coordination of the PBM peptide. Both residues show only three interactions with van der Waals contact distance to residues of 3PDZ1 and 3PDZ4 domain (Fig. S2, S3). Yet, the longer PBM peptide results in a complete hydrogen-bond pattern between the two anti-parallel β-strands formed by the last 5 residues of the URAT1 PBM peptide and β2 of the PDZ domain (Fig. S2, S3). Whereas the binding modes of F^0^, T^-2^, S^-3^ of the 7-mer and 5-mer URAT1 PBM peptide are nearly identical, some differences are noted for the coordination of the Q^-1^ residue in both structures and the K^-4^ in the 3PDZ1^VLKSTQF^ structure (Fig. S2, S3). In the structures 3PDZ1^VLKSTQF^ and 3PDZ4^VLKSTQF^, the Q^-1^ residue forms only water-mediated H-bonds to the PDZ domains instead of direct H-bonds as in 3PDZ1^KSTQF^ and 3PDZ4^KSTQF^ structures (Fig. S2, S3). Notably, in the structure 3PDZ1^VLKSTQF^, the K^-4^ residue side chain reorients and points towards helix α2 as in the structures of the other domains with URAT1 PBM peptide in this study (3PDZ2^KSTQF,^ 3PDZ4^KSTQF^, 3PDZ4^VLKSTQF^ 1PDZ1^KSTQF^) resulting in a second main-chain to main-chain H-bond.

### The interaction of PDZK1 PDZ domains with URAT1 is characterized by fast kinetics

We next analyzed kinetics of the interaction between all PDZ domains of PDZK1 with a URAT1 peptide using surface plasmon resonance (SPR). The biotinylated URAT1 peptide was immobilized on the chip surface, and PDZ domains were flushed over the chip. We observed a significant change of the response for 3PDZ1, 3PDZ2, and 3PDZ4 that indicates binding of these domains to the immobilized URAT1 peptide (Fig. S4A, B, D). The response for 3PDZ3 was very weak even at the highest concentration used (Fig. S4C) consistent with the very low affinity observed in the FA assay (Table 1). We could fit the response curves for 3PDZ1 and 3PDZ4 domains and obtained respective association and dissociation rate constants, k_a_ and k_d_ (Fig. S4A, D; Table 2). The 3PDZ1 domain has a slightly higher on-rate (*k*_a_ = 4.48 ± 0.04 ×10^5^ M^-1^s^-1^), and a slightly lower off-rate (*k*_d_ = 0.4938 ± 0.0038 s^-1^) as compared to the 3PDZ4 domain (*k*_a_ = 3.54 ± 0.01 ×10^5^ M^-1^s^-1^ and *k*_d_ = 0.6306 ± 0.0025 s^-1^), and calculated *K*_D_ values are 1.10 µM and 1.78 µM, respectively (Table 2). We also determined *K*_D_ values from the steady-state response for three out of four analyzed PDZ domains (Fig. S4E, F). These values were very close to the values calculated from kinetic parameters for 3PDZ1 and 3PDZ4 domains. *K*_D_ values for the interaction of PDZ domains with the URAT1 peptide determined here with SPR for the individual domains are 1.3 – 7.4-fold higher than those from the FA assay (see Table 1, Table 2 and Fig. S4F). This might be due to a lower accessibility of the immobilized peptide in SPR as compared to the free peptide in solution in the FA assay. Yet consistently, SPR analysis shows also a higher affinity for 3PDZ1 over 3PDZ4. Taken together, the SPR analysis showed that the high-affinity interaction between URAT1 and PDZK1 in the micromolar range is highly dynamic with high on- and off-rates.

**Table 2.** Kinetic parameters of 3PDZ1 and 3PDZ4 domain binding to the URAT1 peptide determined with SPR.

| | $k_a$ ($\text{M}^{-1}\text{s}^{-1}$) | SE ($k_a$) | $k_d$ ($\text{s}^{-1}$) | SE ($k_d$) | $K_D$ ($\mu\text{M}$) |
| --- | --- | --- | --- | --- | --- |
| 3PDZ1 | 4.48E+05 | 3.50E+03 | 0.4938 | 0.0038 | 1.10 |
| 3PDZ4 | 3.54E+05 | 1.40E+03 | 0.6306 | 0.0025 | 1.78 |

### PDZK1 can bind two URAT1 peptides simultaneously

Considering on one side, the high affinities of the 3PDZ1 and 3PDZ4 domains for URAT1 and, on the other side, the high association and dissociation rates of these interactions, we performed analytical gel filtration to gain information on the stability and stoichiometry of PDZK1:URAT1 complexes. We formed complexes of full-length PDZK1 with the URAT1 peptide fused C-terminally to GFP (GFP^URAT1^) and analyzed them with gel filtration monitoring absorbance at 280 nm and 490 nm (Fig. 6A). This allows quantification of the concentration of both proteins along the elution profiles (Fig. 6B). When GFP^URAT1^ and PDZK1 proteins were combined, they eluted substantially earlier than the individual proteins, indicating the complex formation (Fig. 6A). When they were mixed at 1:1 ratio, both proteins were coeluting in the same peak at near equal concentrations (5.1 µM PDZK1, 6.2 µM GFP^URAT1^ at peak maximum) (Fig. 6B, top). In addition, small amounts of free PDZK1 and free GFP^URAT1^ were detected. Upon increase of GFP^URAT1^ concentration in the sample, more of the protein co-eluted with PDZK1 and no free PDZK1 was observed. In addition, the peak for the complex shifted to even lower elution volume (Fig. 6B). In the sample with 2:1 and 3:1 GFP^URAT1^:PDZK1 ratios, the concentration of GFP^URAT1^ at the maximum of the peak of the complex was 1.4 and 1.5-fold higher than that of PDZK1, respectively. This implies that a mixture of 1:1 and 2:1 complexes was present in this peak in all analysed samples, with more 2:1 complexes at higher GFP^URAT1^ concentration. Even at a concentration substantially higher than the *K*_D_ values of 3PDZ1 and 3PDZ4 domains for the URAT1 peptide, a full saturation of both binding sites was not reached under the experimental conditions, in line with high dissociation rates determined above by SPR (Fig. 6B, bottom).

**Figure 6.**
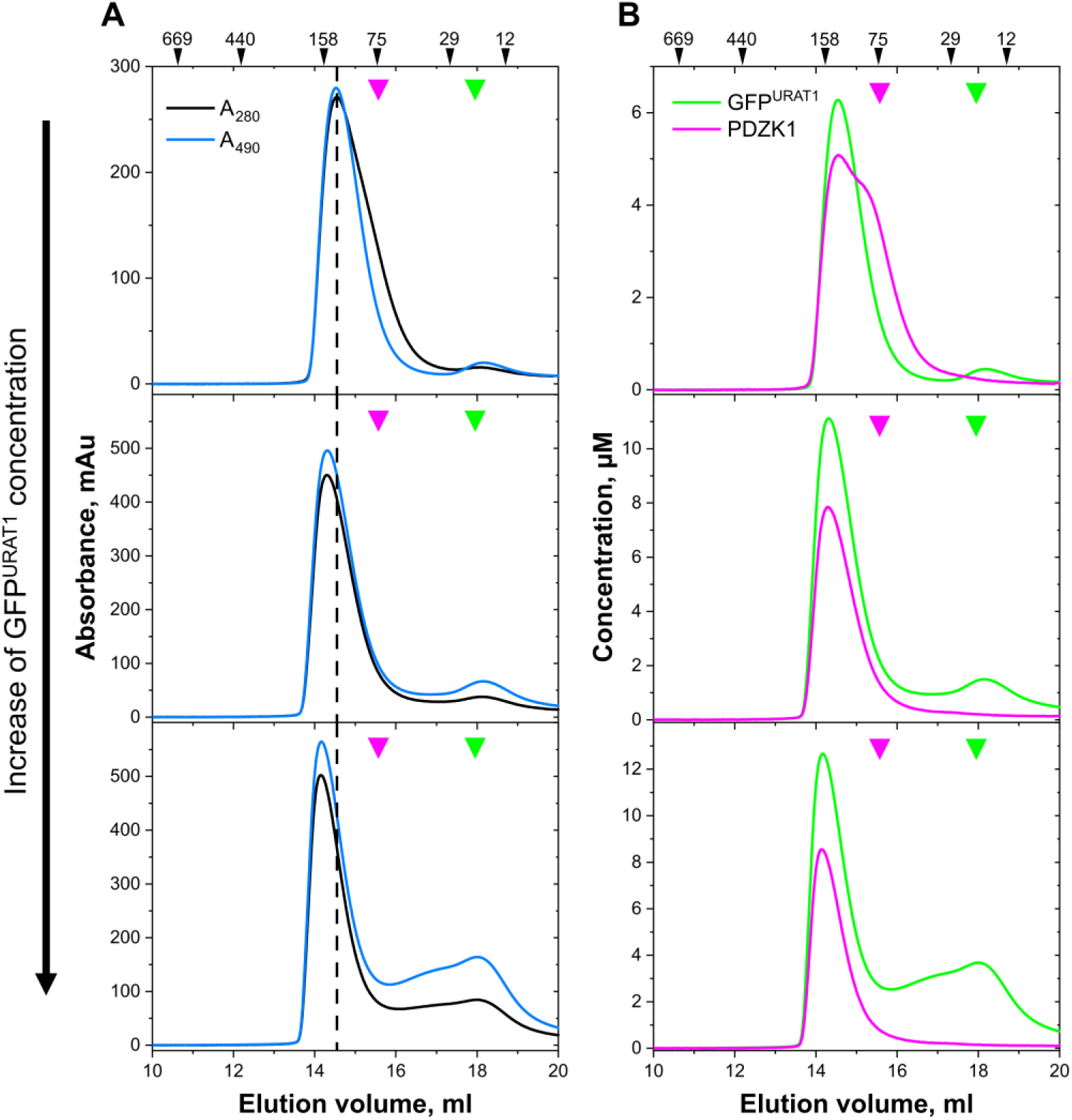
PDZK1 forms dynamic complexes simultaneously binding at least two URAT1 peptides. (**A**) Analytical gel filtration profiles of samples containing 20 µM of full-length PDZK1 and 20, 40, or 60 µM (from top to bottom) of GFP^URAT1^. Absorbance at 280 nm (A_280_) and 490 nm (A_490_) was monitored. Positions of elution maxima of PDZK1 and GFP^URAT1^ alone are marked with large magenta and green triangles, respectively. Black dashed line indicates the maximum of elution profile of the sample containing an equimolar ratio of PDZK1 and GFP^URAT1^, highlighting the shift of the peak with increasing GFP^URAT1^ concentration. Positions of standard proteins used for calibration and their respective molecular mass in kDa are indicated on top of chromatograms. (**B**) Concentration of each protein along the gel filtration profiles quantified from A_280_ and A_490_ as described in Materials and Methods. The concentration of GFP^URAT1^ in the peak corresponded to the complex exceeds that of PDZK1, and the PDZK1/GFP^URAT1^ ratio reaches about 1.5 for the sample with the highest GFP^URAT1^ concentration (bottom).

### Murine Nherf1 binds Urat1 with submicromolar affinity in contrast to the human orthologs

Above we found that, unlike PDZK1, human NHERF1 binds only weakly to URAT1 (Fig. 2), although the latter interaction has been shown for the murine orthologs (Cunningham et al., 2007, Gisler et al., 2003). We hypothesized that this discrepancy might derive from the different PBMs of human URAT1 (KSTQF) and murine Urat1 (MSTRL) (Fig. 7A), as the residues of PDZK1 and NHERF1 PDZ domains involved in the PBM interaction are highly conserved between both species (Fig. 7B, C). We produced and purified full-length mouse Pdzk1 and Nherf1 (Fig. S1B) and determined their *K*_D_^app^ for the peptide that corresponds to 15 C-terminal residues of murine Urat1 (‘Urat1 peptide’) and for NaPi-IIa peptide (Fig. 7D, E). The latter was included for comparison, as the 15 C-terminal residues and particularly the PBM (NATRL) of NaPi-IIa are identical in human and mouse.

**Figure 7.**
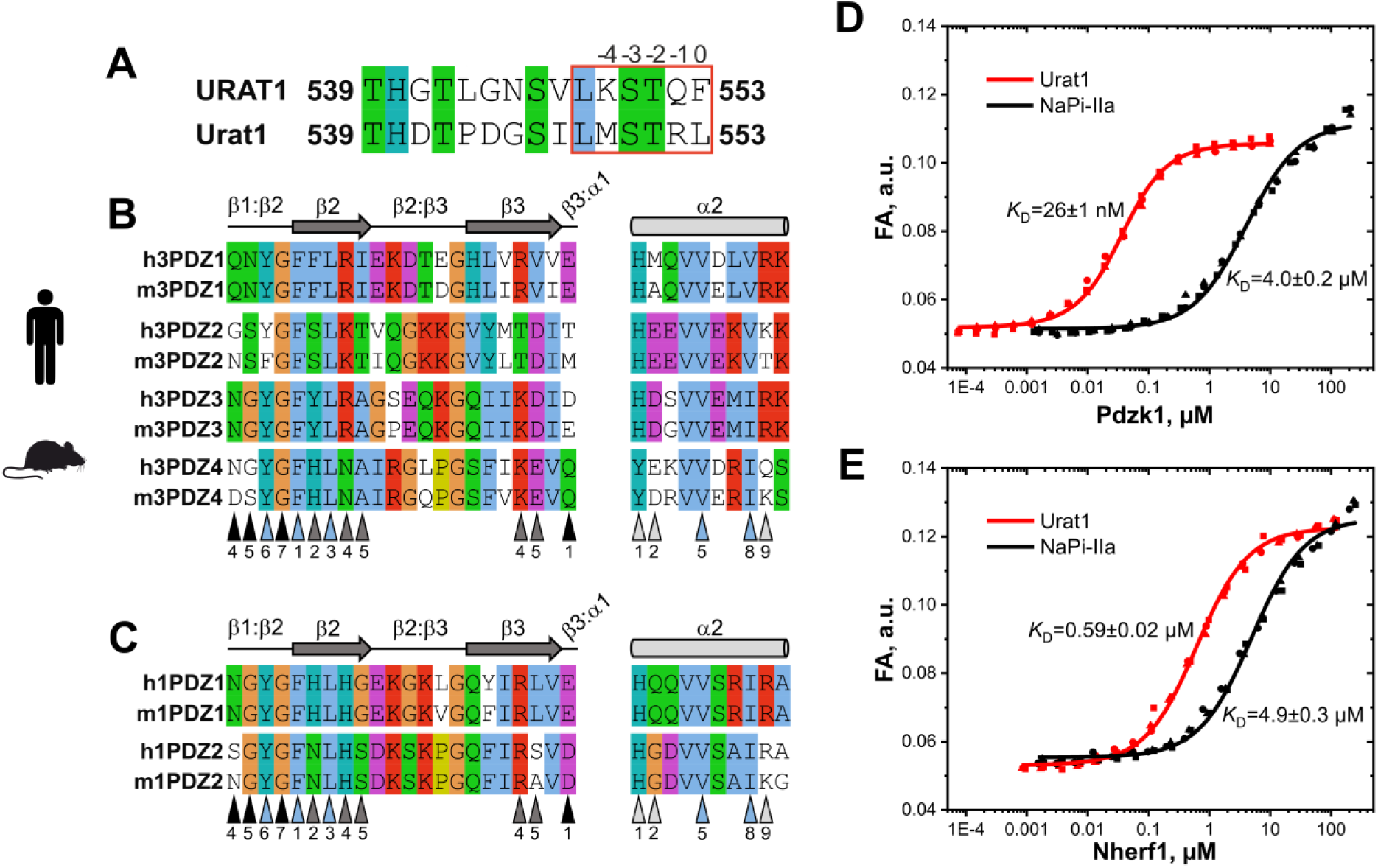
Both mouse Pdzk1 and Nherf1 bind Urat1 peptide with a high affinity. Pairwise alignments of C-terminal sequences of human URAT1 and mouse Urat1 (**A**), and PDZ domain regions of human (h) and mouse (m) PDZK1 (**B**) and NHERF1 (**C**) involved in the interaction with the URAT1 PBM peptide as revealed in the present study (Fig. 4, 5). PBM is outlined in (**A**) with a red rectangle, residue positions are indicated on top. Annotation in (**B**) and (**C**) is as in Fig. 4D. Note, the high similarity of key residues between mouse and human PDZ domains. Binding affinities (*K*_D_^app^ values) of Pdzk1 (**D**) and Nherf1 (**E**) for the fluorescently labeled Urat1 (black) and NaPi-IIa (red) peptides determined with the FA assay. Each measurement was done with three independent biological replicates (data points of different shape) and a concatenated fit was performed (solid lines) for *K*_D_^app^ determination. Respective *K*_D_^app^ ± SE values indicated as “*K*_D_” are shown.

The murine Urat1 peptide binds to Pdzk1 with high affinity (Fig. 7D). The *K*_D_^app^ value was even lower as compared to the human orthologs (26 ± 1 nM for Pdzk1/Urat1 *vs.* 173 ± 6 nM for human PDZK1/URAT1). In contrast, the interaction between NHERF1 and URAT1 is clearly species-specific. We observed an affinity for the interaction between NHERF1 and the Urat1 peptide that was more than 100 times higher than that of the human orthologues (*K*_D_^app^ 0.59 ± 0.02 µM, Fig. 7E vs. >70 µM, see Fig. 2B). As expected from the high similarity of PDZ domains of human and murine PDZK1 and NHERF1 (Fig. 7B, C), their affinities for the NaPi-IIa peptide are very similar. *K*_D_^app^ values of these interactions differ only two-fold between PDZK1 and Pdzk1 (8.1 ± 0.5 µM and 4.0 ± 0.2 µM), and are very close (4.0 ± 0.2 µM and 4.9 ± 0.3 µM) for NHERF1 and Nherf1 (see red curves in Fig. 2 and Fig. 7D, E).

Thus, binding of mouse Nherf1 to Urat1 peptide is another example for high-affinity PDZ domain/PBM interaction. As a control, we determined that both human and mouse NHERF1/Nherf1 bind NaPi-IIa peptide with the micromolar affinity. NaPi-IIa has a C-terminal TRL motif identical to Urat1. The same motif is present in human cystic fibrosis transmembrane conductance regulator (CFTR) that was described to bind with nanomolar affinity to 1PDZ1 domain (Cushing et al., 2008, Wang et al., 1998). We determined the structure of the human 1PDZ1 in complex with NaPi-IIa PBM peptide (1PDZ1^NATRL^, Fig. S5) and revealed a similar binding mode as described previously for the CFTR PBM peptide (1PDZ1^QDTRL^ structure) (Karthikeyan et al., 2001). R^-1^ plays a central role in the specific interaction between 1PDZ1 domain and the bound PBM peptide in both cases. The side chain of this residue forms a salt bridge with the carboxyl of E(β3:α1-1) and an H-bond with main chain oxygen atom of N(β1:β2-5). At the same time, 1PDZ1^NATRL^ lacks ion pairs formed by D^-3^ with R(β3-4) and H(β2-2) present in 1PDZ1^QDTRL^ (Fig. S5). Mouse Urat1 has a serin residue at the −3 position, which most likely forms H-bonds with R(β3-4) and H(β2-2) as we observed in the 1PDZ1^KSTQF^ structure (see Fig. 4F). The H-bonds can explain why the affinity of Nherf1 to the Urat1 peptide is higher than that to the NaPi-IIa peptide (see Fig. 7) and lower than the affinity between human NHERF1 and CFTR peptide (Cushing et al., 2008).

Together, our data indicate a preferential binding of PDZK1 to the URAT1 transporter with nanomolar affinity over NaPi-IIa with micromolar affinity in both mouse and human. In contrast, NHERF1 preferentially binds NaPi-IIa but not URAT1 in human, whereas it binds both transporters in mouse.

## DISCUSSION

Both PDZK1 and NHERF1, multi-PDZ domain scaffold proteins, are highly abundant in proximal tubule cells and are predominantly localized at the brush border membrane, where renal SLCs mediate essential excretion and reabsorption of ions and metabolites (Custer et al., 1994, Enomoto et al., 2002, Gisler et al., 2001, Wade et al., 2001). These two scaffold proteins can bind SLCs via their four and two PDZ domains, respectively (Brone & Eggermont, 2005, Gisler et al., 2003, Walsh et al., 2015). This includes the regulatory interaction with URAT1, a key renal transporter involved in urate reabsorption. Both PDZK1 and NHERF1 have been shown to bind and regulate its proper localization and function in the proximal tubule, although the molecular basis of these interactions is not well understood and existing data for NHERF1 were limited to murine proteins (Anzai et al., 2004, Cunningham et al., 2007, Gisler et al., 2003, Walsh et al., 2015). Scaffold proteins have also been proposed to bind multiple transporters simultaneously to coordinate their function, thereby optimizing secretion and absorption processes in the proximal tubule (Anzai et al., 2007, Brone & Eggermont, 2005, Walsh et al., 2015). A specific model of such organization, the ‘urate transportosome’, involving PDZK1 and several urate transporters including URAT1 was suggested to support urate transport (Anzai et al., 2007, Leask et al., 2024, Prestin et al., 2017, Wright et al., 2010). High-affinity and domain-specific binding of transporters to distinct PDZ domains may promote the spatial organization of such assemblies. Yet, direct experimental evidence for the physical association of different transporters into such a complex is lacking.

We have now clearly shown that full-length human PDZK1, but not NHERF1, binds to the C-terminal PDZ-binding motif of human URAT1 with nanomolar affinity. Such a high affinity is ideally suited to form stable PDZK1/URAT1 complexes in the cell. In contrast, with the very low affinity (*K*_D_^app^>70 µM) determined for the binding of full-length NHERF1 to URAT1, this interaction may be transient or not physiologically relevant. In order to find out whether affinity is based on interaction with single or multiple domains, we compared affinities of all individual PDZ domains of PDZK1 and NHERF1 to respective affinities of the full-length protein using an FA assay. We observed that the 3PDZ1 domain alone binds URAT1 with *K*_D_ of 0.16 ± 0.01 µM, as strongly as the full-length PDZK1 (*K*_D_ = 0.17 ± 0.01 µM). In general, the affinity of PDZ proteins for target proteins was typically obtained with isolated PDZ domains or with PDZ domains fused to large carrier proteins (Amacher et al., 2020, Gogl et al., 2022). Comparison of the affinities of the full-length PDZK1 and those of its individual domains allowed us to exclude allosteric effects on the interaction of high-affinity binding site (3PDZ1) with URAT1 peptide in the full-length protein. Interestingly, in a single study, where a similar approach was applied for PDZK1, negative and positive allosteric effects were observed for interaction of PDZK1 with NHE3 (SLC9A3) and PEPT2 (SLC15A2) peptides, respectively (Hajizadeh et al., 2018). Thus, allosteric effects might be specific for the target protein. The affinity of PDZK1 for URAT1 provided by the 3PDZ1 domain is comparable to that of the strongest PDZ domain/PBM interactions described so far using solution based assays, including a *K*_D_ of 70 nM and 232 nM measured for PDZ2 domain of NHERF2 and the CFTR peptide (Amacher et al., 2013, Cushing et al., 2008), 116 nM for the PDZ1 domain of NHERF1 and the CFTR peptide (Cushing et al., 2008), 150 nM for the PDZ3 domain of PSD95 with the CRIPT peptide (Chi et al., 2012), 190 nM for TIP-1 and β-catenin peptide (Zhang et al., 2008). In comparison to 3PDZ1, the 3PDZ4 domain binds the URAT1 peptide 8-fold weaker, yet the *K*_D_ value is still relatively low (1.35 ± 0.06 µM) indicating that this domain can serve as the secondary binding site for URAT1 and could contribute to the formation of larger complexes. In contrast, 3PDZ2 domain with a determined affinity of 22.1 ± 1.9 µM is likely to contribute substantially less to the interaction of PDZK1 and URAT1, and the URAT1 binding to the 3PDZ3 domain is rather non-specific with an estimated *K*_D_ above 200 µM. Our data are in a good agreement with data from a systems biology approach using a high-throughput holdup assay (Gogl et al., 2022). Gogl et al. have shown that the URAT1 peptide binds to 3PDZ1 with *K*_D_ of 0.68 µM and to 3PDZ4 with *K*_D_ of 3.5 µM; the affinity for the 3PDZ3 domain was below the threshold of 100 µM, and binding to the 3PDZ2 domain was not assessed (Gogl et al., 2022).

Using SPR, we show that the complexes formed by the 3PDZ1 and 3PDZ4 domains with the URAT1 peptide are highly dynamic despite the high affinity. We observed high association and dissociation rates for both domains. Association constants (*k*_a_) in the range 10^5^-10^6^ M^-1^s^-1^ indicate that the complex formation is diffusion-controlled without large rate-limiting conformational changes (Schreiber et al., 2009) as one would expect for PDZ domain/PBM interaction (Amacher et al., 2020, Gianni et al., 2005). Steady-state *K*_D_ values and those calculated from kinetic parameters are very similar, in a low micromolar range. It should be noted that *K*_D_ values for binding of the URAT1 peptide to the PDZ domains of PDZK1 previously determined with SPR were much lower than those obtained with both FA assay and SPR in our study (Anzai et al., 2004). Immobilization of binding partner(s) on a SPR chip surface can influence affinity measurements by surface avidity effects resulting in overestimation of affinity compared to solution-based assays (Amacher et al., 2020, Cushing et al., 2008). In contrast to the previous study (Anzai et al., 2004), we used isolated PDZ domains and immobilized very low amount of the biotinylated URAT1 peptide in our SPR measurements to avoid surface avidity effects. This enabled us to determine the kinetic parameters of PDZ domains with high accuracy resulting in consistent *K*_D_ values obtained with two independent techniques. The SPR data support our conclusion based on the results of the FA assay that 3PDZ1 and 3PDZ4 domains are preferential binding sites for URAT1 in PDZK1, with the interaction being highly dynamic. PDZ domain proteins, and scaffold proteins in general, were considered to be platforms of stably associated proteins. Yet, it has been recognized that many of these proteins can bind their targets dynamically, enabling rapid turnover and the potential for regulation, particularly in signaling pathways (Garbett & Bretscher, 2014). In the case of the PDZK1-URAT1 interaction, one can speculate that such dynamics might serve for fast regulation of the transporter activity in the proximal tubule cells to control the level of urate excretion.

Despite the high relevance of the interactions between PDZ scaffold proteins and many members of the SLC family (Biber et al., 2005, Brone & Eggermont, 2005, Frommelt et al., 2025, Walsh et al., 2015), to our knowledge, only one structure of a PDZ domain in complex with a PBM peptide of a human SLC is available, namely of the PDZ4 domain of PDZK1 with SLC15A2 (PEPT2 transporter) peptide (Hajizadeh et al., 2018). We now expanded this information with seven high-resolution X-ray structures of four PDZ domains of PDZK1 and NHERF1 in complex with the URAT1 and NaPi-IIa PBM peptides contributing to a general understanding of PDZ domain/SLC interactions. The unique set of PDZ domain structures with the same URAT1 PBM peptide bound covers binding affinities ranging from nanomolar to high micromolar *K*_D_ values. The detailed comparative analysis of their binding modes revealed structural features that explain a selectivity of URAT1 for PDZK1 over NHERF1 and, specifically, the higher affinity of the URAT1 peptide for 3PDZ1 and 3PDZ4 domains. In all structures obtained in this study, the canonical PDZ domain/PBM interactions including the β-sheet extension, the carboxylate coordination, the F^0^ binding in the hydrophobic pocket and the coordination of the T^-2^ side chain (Lee & Zheng, 2010, Songyang et al., 1997) are conserved with a few notable exceptions. First, in the 3PDZ2^KSTQF^ structure, corresponding to the medium affinity binding, the peptide’s C-terminal carboxylate moiety is hydrogen-bonded to the side chain oxygen of the S(β1:β2-5) instead of canonical H-bonds with the main chain nitrogen atoms of the carboxylate binding loop. This could contribute to the weaker interaction between 3PDZ2 domain and the PBM peptide. Interestingly, the same non-canonical carboxylate coordination was observed in structures of 3PDZ2^IKITKF^ (Ernst et al., 2014), as well as of the 12^th^ PDZ domain of Multiple PDZ domain protein (MPDZ) with fused ETSV peptide (Elkins et al., 2007). The affinity for the former interaction is not known, but for the latter a very weak affinity with a *K*_D_ of 300 µM was determined (Gogl et al., 2022). The second variation from the canonical binding mode was observed for the coordination of the T^-2^ side chain in the 3PDZ4^KSTQF^ structure, which has a tyrosine residue at α2-1 position instead of histidine. Tyrosine at this position was suggested to be characteristic for class III PDZ domains, which bind PBMs with an acidic residue at position −2, whereas class I PDZ domains are thought to have a histidine residue at α2-1 position and preferentially bind to T^-2^ such as in the URAT1 PBM (Amacher et al., 2020, Hung & Sheng, 2002, Liu & Fuentes, 2019). Based on our data, we can conclude that, not only histidine, but also tyrosine in the α2-1 position enables a strong binding of a class I PBM.

To specifically recognize and bind with a high affinity a defined PBM peptide, additional interactions involving PBM side chains and PDZ domain-specific residues are required. The residues lining the hydrophobic pocket, in which the side chain of F^0^ binds, as well as the residue coordinating the T^-2^ side chain are conserved between PDZ domains analyzed (see Fig. 4D). In contrast, the residues coordinating other PBM residues, Q^-1^, S^-3^, and K^-4^, vary in nature and in position in the PDZ domain (see Fig. 5). 3PDZ1 and 3PDZ4 domains, which bind the URAT1 peptide stronger, have a better coordination of Q^-1^ and S^-3^ side chains in the respective structures. Their side chains form more direct H-bonds compared to the lower affinity complexes with 3PDZ2 and 1PDZ1. The K^-4^ side chain can contribute to the peptide recognition by 3PDZ4 and 3PDZ2 domains by short range electrostatic interactions. As the respective bond cannot be provided by the 3PDZ1 domain, the strongest binder, due to lack of the negatively charged residue at position α2-2, this interaction does not appear to be critical for a high-affinity binding. Taken together, despite the sequence variation of the individual PDZ domains, high-affinity interactions with URAT1 peptide are achieved by multiple direct H-bonds and non-polar interactions with side chains of Q^-1^ and S^-3^, which together have a cumulative effect, and electrostatic interactions can contribute. On the other hand, 3PDZ2 and 1PDZ1 domains that bind the URAT1 peptide weaker lack direct H-bonds to side chains. Instead, structural water molecules bridge the gaps and water-mediated H-bonds are present. Consequently, one can argue that URAT1 has higher selectivity for PDZK1 over NHERF1, and specifically for 3PDZ1 and 3PDZ4 over other analysed domains, since the residues flanking the canonical binding groove in these two domains contribute additional direct contacts with the PBM peptide.

The structures of 3PDZ1 and 3PDZ4 fused with a 7-residue-long URAT1 PBM peptide show in general a highly similar binding mode compared to those with the 5-residue-long PBM peptide including most interactions with side chains. The residues in positions −5 and −6 contribute only a few weak contacts in both structures. Minor differences were observed mostly in the coordination of Q^-1^. In addition, for 3PDZ1, the slighty longer PBM peptide resulted in reorientation of K^-4^ and thus complete coordination of the antiparallel β-strand. Slight variations between the structures at the level of individual amino acid side chains can be explained by the flexible protein recognition model (FPRM) (Grunberg et al., 2004, Teilum et al., 2009). As inferred from the very high association rates determined with SPR, the binding of the URAT1 peptide to the PDZ domain is diffusion-controlled, meaning that the rate-limiting step is the formation of the encounter complex (Teilum et al., 2009). According to FPRM, the next step after formation of the encounter complex is the conformer selection (the recognition complex) followed by conformational changes resulting in the formation of the final complex. These two steps should be notably faster than the association observed with SPR in line with a lack of large conformational changes upon PDZ domain/PBM interaction (Amacher et al., 2020, Gianni et al., 2005). Thus, we can speculate that the energetic barrier between recognition complex(es) and the final complex is low, which is also consistent with a high dissociation rate. Therefore, different complexes can be present in equilibrium in solution, and the X-ray structures provide snapshots of individual conformers. Similarly to our observations, subtle differences in the PBM binding mode in 1PDZ1^TSTTL^ structures obtained from different crystal forms were described earlier (Jiang et al., 2013, Lu et al., 2013). Therefore, this might be a common property of PDZ domains, which bind to their targets with very rapid kinetics. Taking into account that PDZK1 has two high-affinity binding sites for URAT1 with high association and dissociation rates, we analyzed the stability of the PDZK1/URAT1 complex. Furthermore, using the URAT1 peptide fused to the fluorescent protein (GFP^URAT1^), we could determine the stoichiometry of this complex. Indeed, PDZK1 can bind at least two URAT1 peptides simultaneously. However, we could reach only 1.5/1 ratio of GFP^URAT1^/PDZK1 in the complex even upon a high excess of GFP^URAT1^ in the sample. This is most likely due to the fast kinetics of the interaction, when the second binding site cannot be fully saturated under gel filtration conditions. This observation proves that PDZK can provide a scaffold for target proteins, enabling their association (in our case, two URAT1 molecules) in one complex and the formation of a ‘transportosome’ in a cell, though this complex might be transient.

PDZ3 domain of mouse and rat Pdzk1 was shown to bind the sodium/phosphate transporter NaPi-IIa in yeast two-hybrid assays (Gisler et al., 2003, Gisler et al., 2001) and in pull-down assays from opossum kidney cells (Hernando et al., 2002). In our study, we found that this is the only domain that does not contribute to the interaction between human PDZK1 and URAT1. This could enable NaPi-IIa and URAT1 to bind to PDZK1 simultaneously as they have different domain specificity. Unlike mouse proteins, we revealed that the NaPi-IIa peptide binds only to the PDZ1 domain of human PDZK1. Similar to URAT1, the affinity of this domain for the NaPi-IIa peptide is close to that of the full-length PDZK1 indicating that 3PDZ1 is the primary binding site. The other three domains bind the NaPi-IIa peptide either very weakly or not at all. The binding of both transporters to the same PDZ domain renders the formation of a ternary complex of PDZK1, URAT1, and NaPi-IIa in human cells unlikely. The 50-fold higher affinity of PDZK1 for URAT1 as compared to NaPi-IIa makes URAT1 a preferential binding partner of PDZK1 over NaPi-IIa. This might explain an association of genetic variants in *PDZK1* with serum urate concentrations (Köttgen et al., 2013, Tin et al., 2018), but not with effects in phosphate metabolism. Notably, knock-out of *Pdzk1* in mouse does not affect distribution of NaPi-IIa and several other binding partners in the proximal tubular epithelial cells under normal conditions (Capuano et al., 2005, Kocher et al., 2003). Under phosphate-rich and -low diet, *Pdzk1* knock-out showed an effect on NaPi-IIa abundance on the apical membrane (Capuano et al., 2005, Giral et al., 2011, Hernando et al., 2021, Levi et al., 2019).

Interestingly, *Nherf1* knock-out in mice reduced Urat1 expression in the brush border membrane of proximal tubule cells and increased urinary uric acid secretion (Cunningham et al., 2007, Weinman et al., 2006). Indeed, mouse Nherf1 along with Pdzk1 was described to bind Urat1 (Cunningham et al., 2007, Gisler et al., 2003), however we did not observe a strong interaction between respective full-length human proteins. In line with our results, patients carrying *NHERF1* mutations have only abnormal phosphate reabsorption in kidney (increased TmP/GFR values) and no other proximal-tubule-related dysfunction (Courbebaisse et al., 2012, Karim et al., 2008). We noted that mouse Urat1 bears a different C-terminal PBM, namely residues MSTRL. This PBM is similar to the PBMs of well-known NHERF1 targets NaPi-IIa (NATRL) and CFTR (QDTRL) (Cushing et al., 2008, Mamonova et al., 2015). Indeed, we found that the affinity between mouse Nherf1 and Urat1 peptide is in the submicromolar range, more than 100-fold higher than that of human counterparts. Unlike Nherf1, the PDZK1/URAT1 interaction is similarly strong in mice and humans, despite the different PBMs. The species-specific difference in NHERF1/URAT1 affinity explains effects of *Nherf1* knock-out on urate metabolism in mouse and a lack of similar effects in patients with *NHERF1* mutations. A number of mouse models have been developed to study human hyperuricemia (Lu et al., 2019). Notably, serum urate concentration is naturally much higher in higher primates than in mice due to the absence of active uricase, and thus it requires a species-specific regulation (Lu et al., 2019). The species-specific interaction pattern between URAT1 and the two scaffold proteins described in this study contributes to understanding the molecular basis of the different regulation of urate homeostasis in mice and humans.

## MATERIALS AND METHODS

### Protein production and purification

*E. coli* codon-optimized coding sequences of full-length human and mouse PDZK1/Pdzk1 and NHERF1/Nherf1 and of individual PDZ domains of the human proteins in vector pET30a(+) between NdeI and XhoI sites were obtained from BioCat GmbH (Heidelberg, Germany). PDZ domain borders were defined based on previously determined structures and on sequence alignment of different PDZ domains (see Fig. 4D). All constructs used in this study and their properties are listed in Table S1. For production of full-length PDZK1 proteins with N-terminal His_8_-tag, *E. coli* BL21(DE3) cells were transformed with the corresponding plasmid and seeded on LB-agar plates containing 50 µg/ml kanamycin (LB^kana^-agar). Next day, a single colony was inoculated in LB^kana^ and incubated at 37⁰C overnight. Precultures were added to a larger volume of LB^kana^ at 1:50 ratio, and main cultures were incubated at 37⁰C until OD_600_ reached 0.6, when expression was induced with 0.5 mM IPTG. After induction, *E. coli* cultures were incubated at 25⁰C for 5 h and harvested at 10,000g. Production of full-length His_8_-tagged NHERF1 proteins and all individual PDZ domains was performed the same way, except that the expression was induced with 0.1 mM IPTG.

Individual PDZ domains with C-terminal His_8_-tag (all except 3PDZ1, see Table S1) were purified as follows. Cell pellets were resuspended in lysis buffer (50 mM HEPES, 150 mM NaCl, 2 mM EDTA, pH 8.0, 2 mM DTT, 1 mM Pefabloc (Carl Roth GmbH, Karlsruhe, Germany)) and cells were lysed with a cell disruptor (Constant Systems Ltd., Daventry, UK).

Cell debris was removed by centrifugation at 30,000g for 30 min. Cleared cell lysate was loaded on a HisTrap FF 5 ml column (Cytiva, Marlborough, MS, USA) equilibrated with IMAC buffer (20 mM HEPES, 300 mM NaCl, 20 mM imidazol, 1 mM DTT, pH 8.0) and unbound material was washed out with 10 column volumes (CV) of the same buffer. Protein was eluted with a linear imidazole gradient of 20 mM to 300 mM over 10 CV. Fractions containing protein of interest were combined and concentrated to a volume of about 5 ml before loading on HiLoad Superdex75 26/600 column (Cytiva) equilibrated with SEC buffer (20 mM HEPES, 150 mM NaCl, 0.5 mM TCEP, pH 7.5). Eluted fractions were analyzed with SDS-PAGE, and those containing pure protein of interest were combined, concentrated, frozen in liquid nitrogen and stored at −80⁰C. Purification of full-length PDZK1 and NHERF1 proteins was done following the same protocol and using a HiLoad Superdex200 26/600 (Cytiva) instead of the HiLoad Superdex75 26/600 column. For storage, full-length proteins were supplemented with 20% glycerol before freezing.

For purification of tag-less 3PDZ1, as well as of constructs used for crystallization, the N-terminal affinity tag was cleaved with TEV protease. Cell lysis and IMAC were performed as above. For cleavage, fractions after IMAC were mixed with TEV protease (about 1:10 mass ratio) and dialyzed overnight at room temperature against 20 mM Tris-HCl, 150 mM NaCl, 1 mM DTT, pH 8.0. After dialysis, a fresh aliquot of TEV protease was added and the protein was incubated for 1-2 days at room temperature to increase the yield of the cleaved product. Precipitated proteins were removed by centrifugation, and a second IMAC was performed collecting unbound proteins. The gel filtration was performed as above for tag-less 3PDZ1 or using protein buffer for crystallization as described in Table S2. TEV protease was produced and purified according to Kapust et al. (Kapust et al., 2001) with plasmid pRK793 (a gift from David Waugh, Addgene plasmid #8827).

The plasmid encoding the fusion protein consisting of enhanced green fluorescent protein and 15 C-terminal residues of URAT1 (GFP^URAT1^) with N-terminal His_8_-tag in pET30a(+) vector was obtained from BioCat GmbH. The protein production was performed as described above for the full-length PDZK1. The purification of GFP^URAT1^ followed the same protocol as for the His-tagged individual PDZ domains.

The purity of purified proteins was analyzed with SDS-PAGE on 12% (individual PDZ domains) or 4-12% (full-length PDZK1 and NHERF1) NuPAGE Bis-Tris gels (Thermo Fischer Scientific, Waltham, USA).

### Fluorescence anisotropy assay

We performed fluorescence anisotropy (FA) assays using synthetic peptides that correspond to the 15 C-terminal residues of human URAT1 (URAT1 peptide, THGTLGNSVLKSTQF), murine Urat1 (Urat1 peptide, THDTPDGSILMSTRL) or NaPi-IIa (NaPi-IIa peptide, SPRLALPAHHNATRL, identical in mouse and human), which are N-terminally labeled with tetramethylrhodamine (TAMRA). All peptides used in the present study were ordered from BioCat GmbH. To determine affinities, full-length proteins and individual domains were prepared in FA buffer (20 mM HEPES, 150 mM NaCl, 0.5 mM TCEP, pH 7.5) and titrated 1:1 over 18 wells in black no-binding F-bottom 96-well plates (Greiner Bio-One, Frickenhausen, Germany). An equal amount of a master mix containing 0.1 µM (0.04 µM in the case of Pdzk1 + Urat1 peptide) of labeled peptide and 0.2% BSA in FA buffer was added to the protein titration series resulting in final concentrations of 0.05 µM or 0.02 µM peptide and 0.1% BSA. After 30 min incubation at room temperature in the dark, parallel and perpendicular polarized fluorescence was measured at Clariostar Plus plate reader (BMG Labtech, Ortenberg, Germany) with 540/20 nm excitation filter and 590/20 nm emission filter at 25⁰C. Gain in two channels was adjusted to get 75 mP fluorescence polarization for the control sample containing a fluorescent peptide and no protein. Background fluorescence was measured in samples without a fluorescent peptide. Fluorescence anisotropy (r) was calculated with the MARS analysis software (BMG Labtech) after subtraction of the background fluorescence. For *K*_D_ determination, fluorescence anisotropy was plotted against protein concentration and the one-site binding model (equation 1) was used to fit the data points.

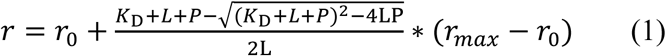

where r_0_ is fluorescence anisotropy in the absence of protein, r_max_ – fluorescence anisotropy of the peptide bound to protein, *K*_D_ – dissociation constant, L – peptide concentration (fixed at 0.05 µM or 0.02 µM), P – protein concentration. All measurements were done with three independently produced and purified protein samples (biological replicates). Data from all replicates were plotted together and a concatenated fit was performed in OriginPro 2020 software (OriginLab Corporation, Northampton, MA, United States). Due to the fixed low concentration of the peptide in the FA assay, only the lowest *K*_D_ value can be determined in the case of proteins having several binding sites, thus we refer to *K*_D_ ^app^ for full-length proteins.

### Protein crystallization

For the crystallization of PDZ domains in complex with PBM peptides of interest, the respective 5- or 7-mers peptides were fused to the C-terminus of the PDZ domains. This method has been successfully used to determine the structures of other PDZ domain/PBM complexes (Birrane et al., 2013, Elkins et al., 2007, Ernst et al., 2014, Jiang et al., 2013, Karthikeyan et al., 2001). Purified proteins were concentrated by spin-filtration (Vivaspin 500 5 MWCO PES Centrifugal Concentrators, Sartorius AG, Göttingen, Germany) and subjected to crystallization at typically 2-3 different concentrations. Crystallization assays were performed using the sitting drop vapor diffusion method in MRC 2-well crystallization plates (Jena Biosciences GmbH, Jena, Germany) and JBScreen Classic HTS I, Classic HTS II and Pi-PEG commercial screens (Jena Biosciences). A Phoenix robot (Art Robbins Instruments, Sunnyvale, CA, USA) was used for pipetting 0.2 µl protein solution with 0.2 µl reservoir solution while the reservoir volume was 75 µl. The conditions in which crystals grew were reproduced and optimized using 0.5 µl of protein and reservoir solutions. Individual crystallization conditions are summarized in Table S2.

### Structure determination and refinement

When required, crystals were cryo-protected by soaking them for 1-5 minutes in a reservoir solution supplemented with cryo-protectant (Table S2). Then, crystals were harvested and flash-cooled in liquid nitrogen. X-ray diffraction data were collected at the European Synchrotron Radiation Facility (ESRF, Grenoble, France) on beamlines MASSIF3 (ID30A-3), ID23-1, ID30B, and BM07 (Table S3 and S4). Diffraction data were processed with AutoProc and STARANISO when data were anisotropic (Vonrhein et al., 2011). The phase problem was solved by molecular replacement using Phaser (McCoy et al., 2007). Search models are listed in Table S2. In all cases, only the PDZ domain part in those search models was used for molecular replacement. Final structures were obtained after several iterations of model building using Coot (Emsley et al., 2010) and refinement using Phenix (Liebschner et al., 2019). The quality of the structures was assessed using MolProbity (Chen et al., 2010). The binding of the PBM peptides was analyzed using the NCONT tool from the CCP4 and a cutoff distance of 4 Å (Winn et al., 2011). Figures were generated with the PyMOL Molecular Graphics System, Version 2.4.0a0 (Schrödinger, LLC; NY, USA).

### Surface plasmon resonance

All experiments were run on a Biacore T200 on a Chip CAP (Cytiva) and at 25 °C using 20 mM HEPES, 150 mM NaCl, 0.5 mM TCEP, 0.05% Tween-20, pH 7.5 as a running buffer. The chip surface was conditioned by injecting the biotin CAPture reagent (Cytiva) at a flowrate of 2 µL/min for 5 min followed by a 25 s injection at 5 µL/min of 0.2 µg/mL N-terminally biotinylated URAT1 peptide (identical in the sequence to one used in the FA assay) in the running buffer. The resulting capture level was 15 RU. For the affinity measurements, a serial dilution with 1:1 buffer:protein steps over 11 points was prepared starting at 10 µM for 3PDZ1, 310 µM for 3PDZ2, 394 µM for 3PDZ3, and 27 µM for 3PDZ4. The interactions were measured as multi-cycle kinetics with a contact time of 120 s and dissociation time of 300 s at a flowrate of 30 µL/min.

Reference and blank subtracted data were analyzed with the Biacore T200 Evaluation software (version 3.0) using the steady-state affinity model or the 1:1 kinetic model with RI jump disabled.

### Analytical gel filtration

To determine the stoichiometry of the complex formed by full-length PDZK1 and URAT1, we used the URAT1 peptide fused to the C-terminus of green fluorescent protein (GFP^URAT1^) and analyzed a complex formation with analytical gel filtration. 20 µM full-length PDZK1 was mixed and GFP^URAT1^ at 1:1, 1:2, and 1:3 molar ratios in FA buffer. 20 µM full-length PDZK1 and 20 µM, 40 µM, or 60 µM GFP^URAT1^ were used as controls. All samples were incubated for at least 30 min at room temperature, loaded on Superdex 200 Increase 10/300 GL column (Cytiva), and eluted with a flow rate of 0.5 ml/min. Protein elution was followed by monitoring absorbance at 280 nm (A_280_) and 490 nm (A_490_). We determined the concentration of each protein along the elution profile in the samples containing both PDZK1 and GFP^URAT1^ as follows. First, we quantified the ratio A_280_/A_490_ for control GFP^URAT1^ samples, which was 0.491. Next, in the elution profiles of samples containing two proteins, we determined the contribution of GFP^URAT1^ to the total A_280_ as A_490_×0.491. To determine the PDZK1 contribution, we subtracted the contribution of GFP^URAT1^ from the total A_280_. Finally, we quantified the concentration (C) of each protein along the elution profile dividing the individual contributions to the total A_280_ by respective molar extinction coefficients at 280 nm (26,360 M^-1^cm^-1^ for PDZK1 and 21,890 M^-1^cm^-1^ for GFP^URAT1^). These calculations are summarized in formulas 2 and 3.

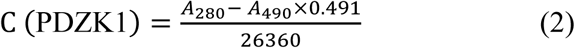

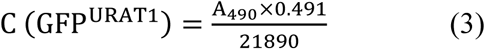

The column was calibrated using thyroglobulin (669 kDa), ferritin (440 kDa), aldolase (158 kDa), conalbumin (75 kDa), carbonic anhydrase (29 kDa), and cytochrome c (12 kDa) as protein standards.

### Use of Generative AI tools

DeepL Write (DeepL SE, Cologne, Germany) and ChatGPT (OpenAI, San Francisco, CA, USA) were used for language editing.

## DATA AVAILABILITY STATEMENT

The data that support the findings of this study are available in the Materials and Methods, Results, and Supplemental Material of this article. Structures will be available in the Protein Data Bank (https://www.rcsb.org/), the accession numbers are provided in Tables S3 and S4.

## AUTHOR CONTRIBUTIONS

Conceived the study - EVM, AK, CH

Prepared proteins - EVM, JIH, JG, CS, AKI

Performed FA assays - EVM, JIH, JG, CS

Crystallized proteins - EVM, CW, JIH, JG, AKI

Performed structural analysis - CW, JIH, BK

Performed and analyzed SPR measurements - EVM, TM

Analyzed the data - EVM, CW, CH

Wrote original draft - EVM, CW, CH

All authors discussed the results and contributed to editing and review of the original manuscript draft.

## ACKNOWLEDGMENTS

This work was funded by the Deutsche Forschungsgemeinschaft (DFG, German Research Foundation) under Germany’s Excellence Strategy (CIBSS – EXC-2189 – Project ID 390939984) in the form of project funding to A.K, T.H., and C.H. and of CIBSS Launchpad Program Funds to E.M.; by the DFG through Project-ID 431984000 – SFB 1453 to A.K. and C.H. and through Project-ID 403222702 – SFB 1381 to C.H. The plasmid pRK793 encoding TEV protease was obtained from Addgene and is a gift from David Waugh. We acknowledge the European Synchrotron Radiation Facility (ESRF) for provision of synchrotron radiation facilities under proposal number MX-2446 and MX-2550. We would like to thank the staff of the ESRF and EMBL Grenoble for assistance and support in using beamlines ID23-1, ID30A-3, ID30B and BM07 under proposal number MX-2446 and MX-2550. We thank Julia Schimpf and Thorsten Hugel (Institute of Physical Chemistry, University of Freiburg) for the introduction to fluorescence anisotropy measurements, and T. Hugel for critical comments on the manuscript.

