## Supplemental information for "Molecular determinants of selective and high-affinity binding of the scaffold protein PDZK1 to the transporter URAT1"

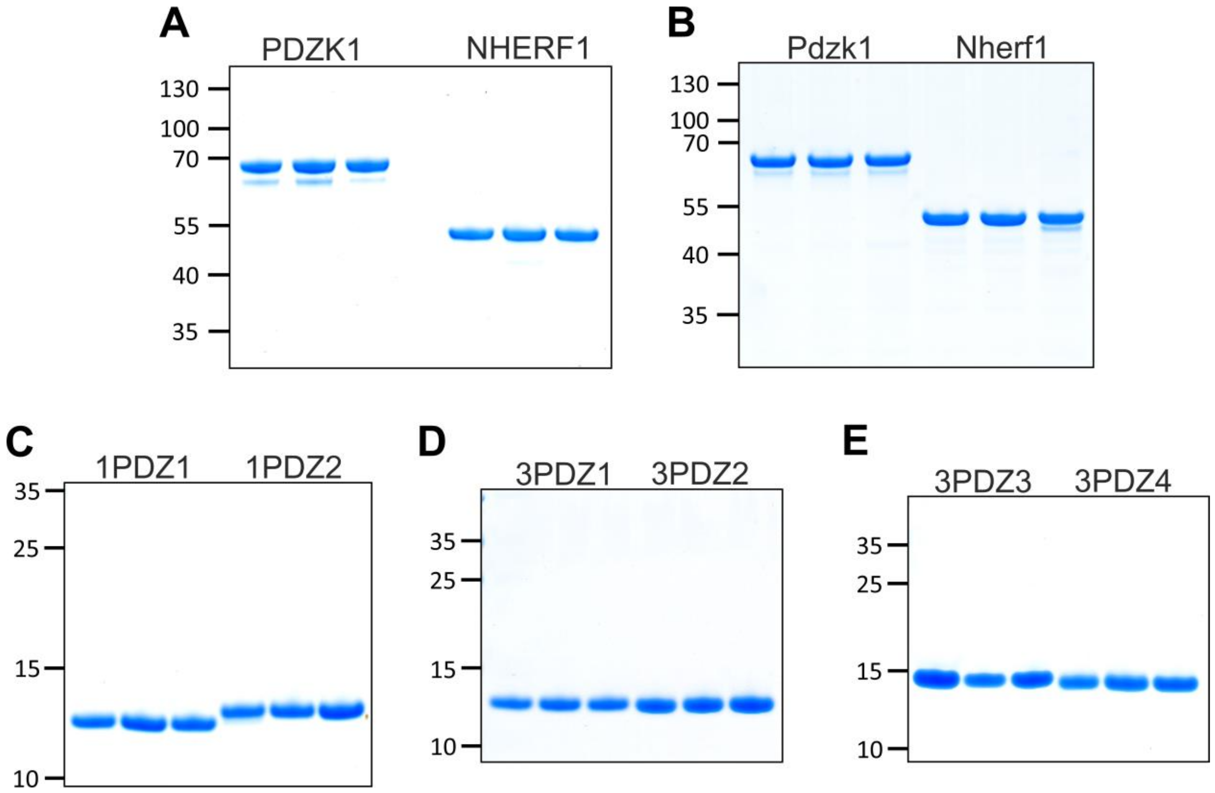

**Fig. S1. Three biological replicates of each protein were independently purified for FA measurements.**

(**A**) Full-length human PDZK1 and NHERF1; (**B**) Full-length mouse Pdzk1 and Nherf1; (**C**) PDZ1 and PDZ2 of human NHERF1; (**D**) PDZ1, PDZ2 and (**E**) PDZ3, PDZ4 of human PDZK1. 1 µg of pure proteins was loaded on 4-12% (**A**, **B**) or 12% (**C-E**) NuPAGE Bis-Tris gels. Positions of co-migrated molecular mass standards are indicated in kDa on the left.

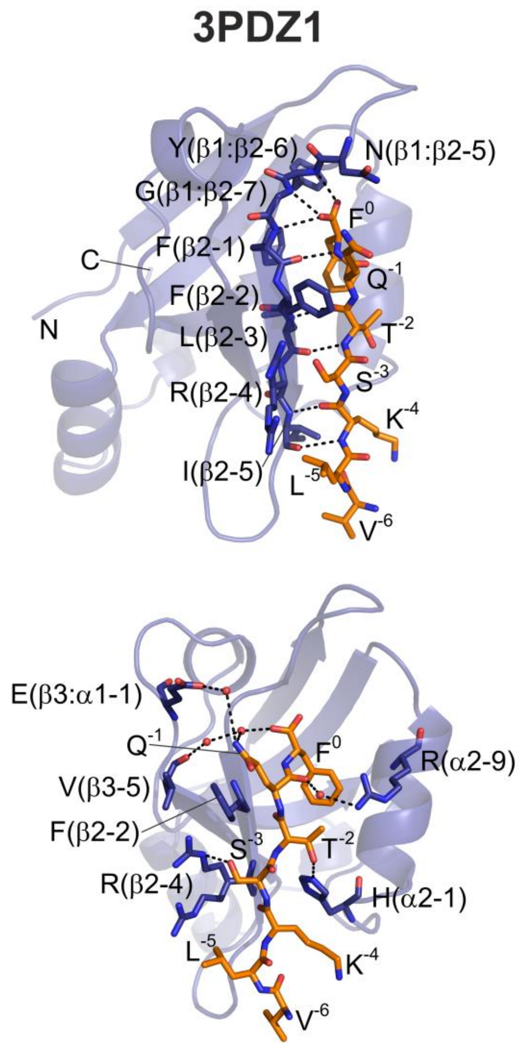

| Peptide | Position in PDZ | 3PDZ1 KSTQF | 3PDZ1 VLKSTQF | Remarks |
|---|---|---|---|---|
| $F^0$ | β1:β2-5 | Asn 19 | Asn 19 | |
| | β1:β2-6 | **Tyr 20** | **Tyr 20** | CBL |
| | β1:β2-6 | Tyr 20 | Tyr 20 | hydrophobic |
| | β1:β2-7 | **Gly 21** | **Gly 21** | CBL |
| | β1:β2-7 | Gly 21 | Gly 21 | hydrophobic |
| | β2-1 | **Phe 22** | **Phe 22** | CBL/β-strand |
| | β2-1 | Phe 22 | Phe 22 | hydrophobic |
| | β2-3 | Leu 24 | Leu 24 | hydrophobic |
| | α2-5 | Val 72 | Val 72 | hydrophobic |
| | α2-8 | Val 75 | Val 75 | hydrophobic |
| | α2-9 | **Arg 76*** | | |
| | α2-9 | Arg 76 | Arg 76 | |
| $Q^{-1}$ | β2-1 | Phe 22 | Phe 22 | |
| | β2-2 | Phe 23 | Phe 23 | |
| | β3:α1-1 | **Glu 39** | **Glu 39** | |
| | α2-9 | **Arg 76*** | **Arg 76*** | |
| $T^{-2}$ | β2-2 | Phe 23 | Phe 23 | |
| | β2-3 | **Leu 24** | **Leu 24** | β-strand |
| | α2-1 | **His 68** | **His 68** | PDZ class |
| | α2-5 | Val 72 | Val 72 | |
| | α2-9 | Arg 76 | Arg 76 | |
| $S^{-3}$ | β2-2 | Phe 23 | Phe 23 | |
| | β2-3 | Leu 24 | Leu 24 | |
| | β2-4 | **Arg 25** | **Arg 25** | |
| | α2-1 | His 68 | | |
| $K^{-4}$ | β2-3 | | Leu 24 | |
| | β2-4 | Arg 25 | Arg 25 | |
| | β2-5 | **Ile 26** | **Ile 26** | β-strand |
| | α2-1 | His 68 | His 68 | |
| $L^{-5}$ | β2-5 | | Ile 26 | |
| $V^{-6}$ | β2-5 | | Ile 26 | |
| | β2:β3-1 | | Lys 28 | |

| | |
|---|---|
| xxx | 2 main-chain to main-chain H-bonds |
| xxx | 1 main-chain to main-chain H-bond |
| xxx | water mediated H-bond |
| xxx | H-bond or salt bridge involving at least one side chain |
| xxx | residues in van der Waals contact distance |

**Fig. S2. The 3PDZ1$^{VLKSTQF}$ structure and interaction analysis.**

On the left, cartoon representation of 3PDZ1$^{VLKSTQF}$ structure (Table S4) as in Figure 4E and 4F. On top left, residues involved in main-chain-to-main-chain H-bonds (black dotted lines) between the 3PDZ1 domain and the URAT1 peptide are shown as sticks. Specific interactions involving the side chains of the PDZ domain residues (shown as sticks) are highlighted as black dotted lines in bottom left panel. Water molecules involved in the coordination of the PBM residues are depicted as red spheres. On the right, comparison of interactions between 3PDZ1 domain and URAT1 peptide in 3PDZ1$^{KSTQF}$ and 3PDZ1$^{VLKSTQF}$ structures with annotation as in Fig. 5.

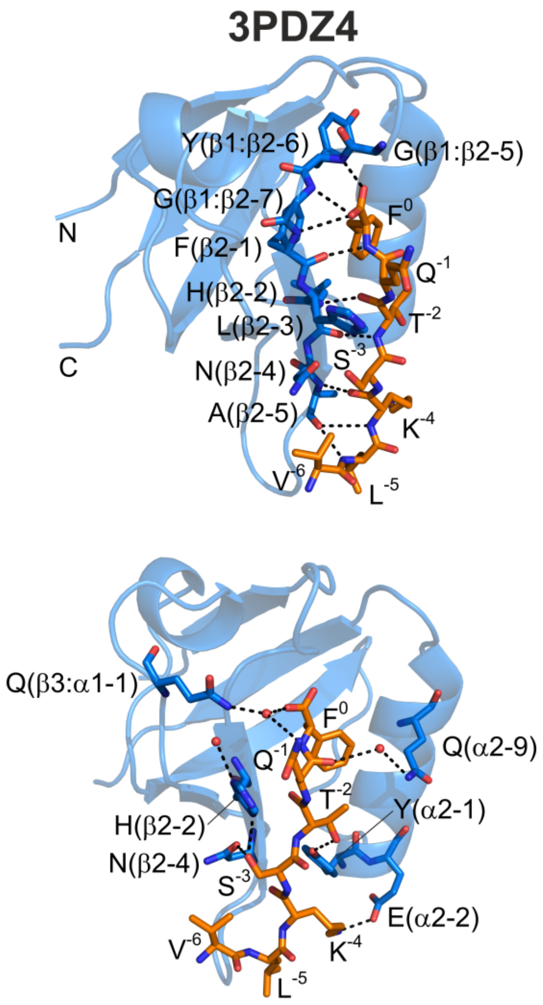

| Peptide | Position in PDZ | 3PDZ4 | | Remarks |
|---|---|---|---|---|
| | | KSTQF | VLKSTQF | |
| F$^{0}$ | β1:β2-4 | **Asn 386** | | |
| | β1:β2-5 | Gly 387 | Gly 387 | |
| | β1:β2-6 | **Tyr 388** | **Tyr 388** | CBL |
| | β1:β2-6 | Tyr 388 | Tyr 388 | hydrophobic |
| | β1:β2-7 | **Gly 389** | **Gly 389** | CBL |
| | β1:β2-7 | Gly 389 | Gly 389 | hydrophobic |
| | β2-1 | **Phe 390** | **Phe 390** | CBL/β-strand |
| | β2-1 | Phe 390 | Phe 390 | hydrophobic |
| | β2-3 | Leu 392 | Leu 392 | hydrophobic |
| | β3:α1-1 | | **Gln 406** | |
| | α2-5 | Val 440 | Val 440 | hydrophobic |
| | α2-8 | Ile 443 | Ile 443 | hydrophobic |
| | α2-9 | Gln 444 | Gln 444 | |
| Q$^{-1}$ | β2-1 | Phe 390 | Phe 390 | |
| | β2-2 | **His 391** | | |
| | β3:α1-1 | | **Gln 406** | |
| | α2-9 | **Gln 444*** | **Gln 444*** | |
| T$^{-2}$ | β2-2 | His 391 | His 391 | |
| | β2-3 | **Leu 392** | **Leu 392** | β-strand |
| | α2-1 | **Tyr 436** | **Tyr 436** | PDZ class |
| S$^{-3}$ | β2-2 | **His 391** | **His 391** | |
| | β2-3 | Leu 392 | Leu 392 | |
| | β2-4 | **Asn 393** | **Asn 393** | |
| | α2-1 | Tyr 436 | Tyr 436 | |
| K$^{-4}$ | β2-3 | | Leu 392 | |
| | β2-4 | Asn 393 | Asn 393 | |
| | β2-5 | **Ala 394** | **Ala 394** | β-strand |
| | β2:β3-2 | Arg 396 | | |
| | α2-1 | Tyr 436 | Tyr 436 | |
| | α2-2 | **Glu 437** | **Glu 437** | |
| L$^{-5}$ | β2-5 | | **Ala 394** | |
| | β2:β3-2 | | Arg 396 | |
| V$^{-6}$ | β2-4 | | Asn 393 | |
| | β2-5 | | Ala 394 | |

| | |
|---|---|
| **xxx** | 2 main-chain to main-chain H-bonds |
| **xxx** | 1 main-chain to main-chain H-bond |
| **xxx** | water mediated H-bond |
| **xxx** | H-bond or salt bridge involving at least one side chain |
| xxx | residues in van der Waals contact distance |

**Fig. S3. The 3PDZ4$^{VLKSTQF}$ structure and interaction analysis.**

On the left, cartoon representation of 3PDZ4$^{VLKSTQF}$ structure (Table S4) as in Figure 4E and 4F. On the right, comparison of interactions between 3PDZ1 domain and URAT1 peptide in 3PDZ4$^{KSTQF}$ and 3PDZ4$^{VLKSTQF}$ structures as in Fig. 5. For details of annotation see legend of Fig. S2.

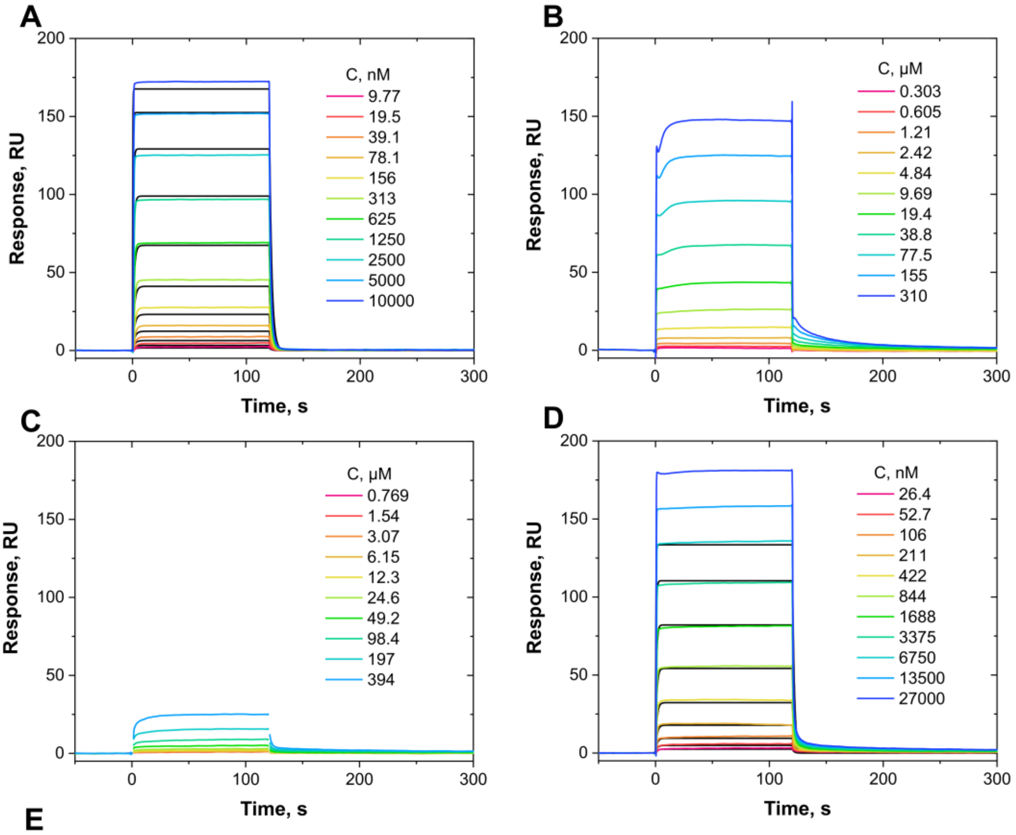

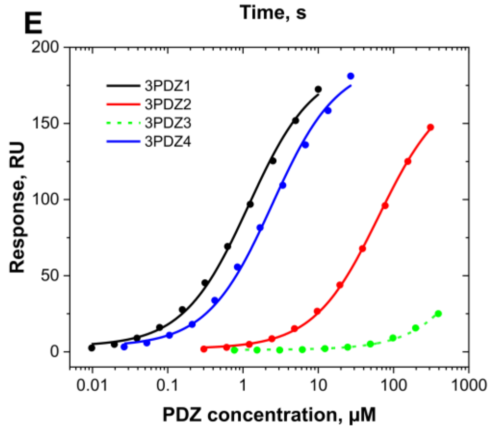

**F**

| Domain | $K_D$ ± SE, µM |
|---|---|
| 3PDZ1 | 1.18 ± 0.08 |
| 3PDZ2 | 63.6 ± 2.2 |
| 3PDZ3 | n.d. |
| 3PDZ4 | 2.45 ± 0.21 |

**Fig. S4. Surface plasmon resonance (SPR) analysis of the interaction between PDZ domains of PDZK1 and the URAT1 peptide.**

(**A-D**) SPR sensogramms for interaction analysis of 3PDZ1 (**A**), 3PDZ2 (**B**), 3PDZ3 (**C**), and 3PDZ4 (**D**) with the immobilized URAT1 peptide. Raw curves corresponding to the samples with different PDZ concentration are in color. The 1:1 kinetic model fit curves for 3PDZ1 (**A**) and 3PDZ4 (**D**) are in black. The kinetic model fit was not possible for the 3PDZ2 and 3PDZ3 domains due to relatively strong non-specific binding of the former in the reference channel and too weak response for the latter. (**E**) Steady-state binding isotherms for four analyzed PDZ domains fitted with one-site binding model, and (**F**) $K_D$ values determined with these isothermes. $K_D$ value of 3PDZ3 could not be determined due to a very weak binding to the URAT1 peptide.

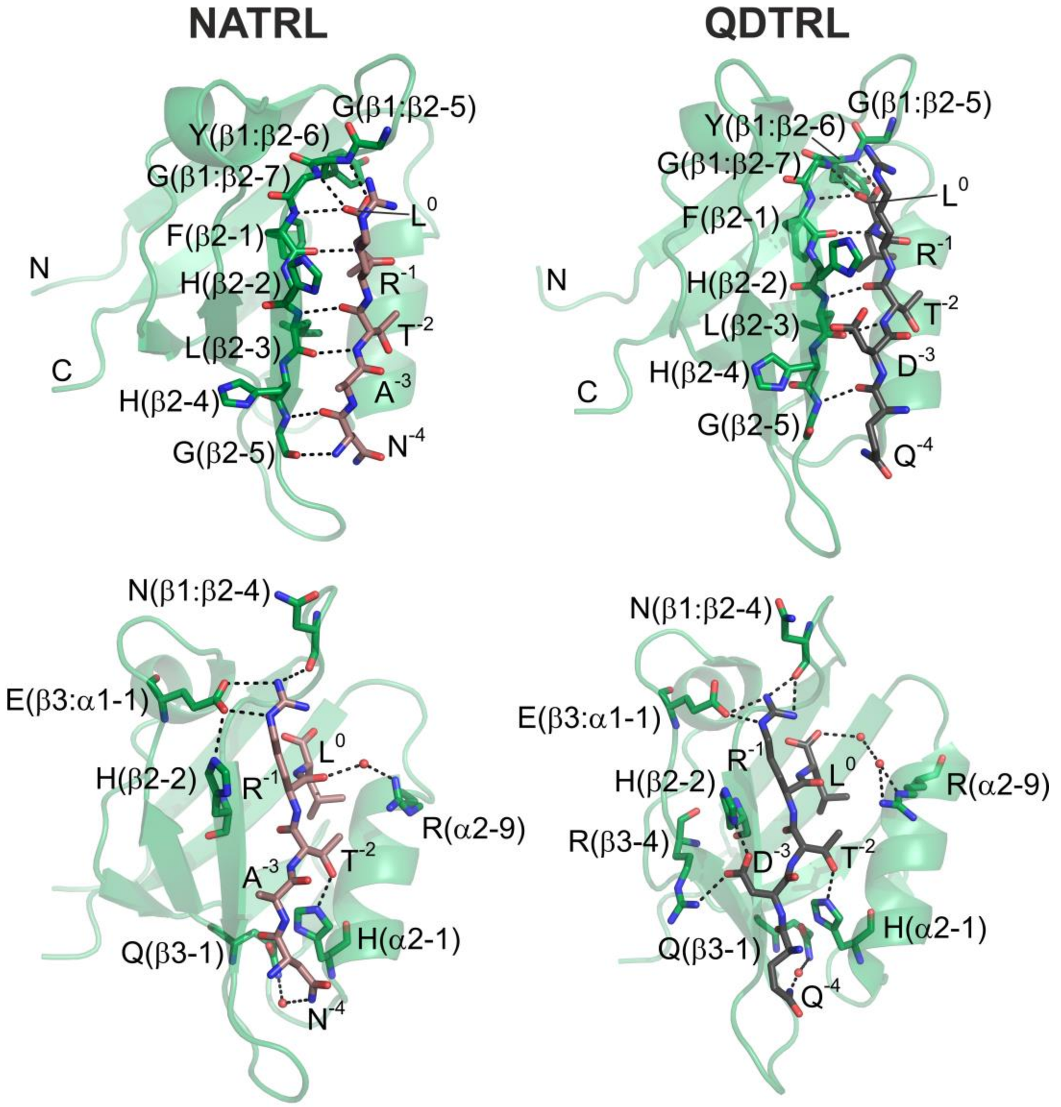

**Fig. S5. The 1PDZ1$^{NATRL}$ structure and comparison with 1PDZ1$^{QDTRL}$ structure.** Representations of 1PDZ1$^{NATRL}$ (Table S4) and 1PDZ1$^{QDTRL}$ (PDB ID 1I92) structures are as in Figure 4E and 4F. NaPi-IIa PBM peptide (NATRL) and CFTR PBM peptide (QDTRL) are shown as brown and grey sticks, respectively.

**Table S1. PDZK1 and NHERF1 constructs used in the study and their properties calculated with Expasy ProtParam tool.**

| Protein | Residues | Mw, kDa | pI | ε280, $M^{-1}cm^{-1}$ |
|---|---|---|---|---|
| **His-PDZK1** | 1-519 | 58.4 | 5.68 | 26360 |
| **3PDZ1** | 6-105 | 11.43 | 6.16 | 2980 |
| **$3PDZ1^{KSTQF}$** | 6-105 | 12.0 | 6.94 | 2980 |
| **$3PDZ1^{VLKSTQF}$** | 6-105 | 12.2 | 6.94 | 2980 |
| **3PDZ2-His** | 131-215 | 10.7 | 6.53 | 4470 |
| **$3PDZ2^{KSTQF}$** | 131-215 | 9.98 | 6.13 | 4470 |
| **3PDZ3-His** | 240-323 | 10.45 | 5.94 | 2980 |
| **3PDZ4-His** | 375-458 | 10.4 | 7.13 | 2980 |
| **$3PDZ4^{KSTQF}$** | 375-458 | 9.7 | 7.96 | 2980 |
| **$3PDZ4^{VLKSTQF}$** | 375-458 | 9.9 | 7.96 | 2980 |
| **His-NHERF1** | 1-357 | 40.1 | 5.99 | 9970 |
| **1PDZ1-His** | 11-94 | 10.1 | 7.13 | 2980 |
| **1PDZ2-His** | 151-234 | 10.45 | 7.93 | 1490 |
| **$1PDZ1^{KSTQF}$** | 11-94 | 9.80 | 8.01 | 2980 |
| **$1PDZ1^{NATRL}$** | 11-94 | 9.77 | 8.02 | 2980 |
| **His-Nherf1** | 1-355 | 39.7 | 6.04 | 8480 |
| **His-Pdzk1** | 1-519 | 57.7 | 5.6 | 17880 |

**Table S2. Crystallization and cryoprotection conditions as well as molecular replacement search models used**

| Protein | Protein buffer condition | Crystallization condition | Cryo protection (crystallization condition supplemented with…) | Structure used as molecular replacement search model |
|---|---|---|---|---|
| $3PDZ1^{KSTQF}$ | 20 mM Tris-HCl, pH 8.0,<br>150 mM NaCl,<br>1 mM TCEP | 50 mM Bicine pH 8.4,<br>12% PEG 600,<br>3.5% PEG 2000 MME | 12.5% ethylene glycol | 4f8k |
| $3PDZ2^{KSTQF}$ | 20 mM Tris-HCl, pH 7.5,<br>200 mM NaCl,<br>0.5 mM TCEP | 26% PEG 1500 | 10% ethylene glycol | 4q2p |
| $3PDZ4^{KSTQF}$ | 20 mM Tris pH 8.5,<br>150 mM NaCl,<br>1 mM TCEP | 1% 2-methyl-2,4-pentanediol,<br>2.5 M ammonium sulfate | 12.5% ethylene glycol | 6ezi |
| $1PDZ1^{KSTQF}$ | 20 mM Tris-HCl, pH 8.0,<br>150 mM NaCl,<br>1 mM TCEP | 100 mM sodium acetate pH 4.8<br>200 mM ammonium acetate<br>24% PEG 4000 | 25% glycerol | 4mpa |
| $3PDZ1^{VLKSTQF}$ | 20 mM Tris-HCl, pH 8.0,<br>150 mM NaCl,<br>1 mM TCEP | 100 mM MES, pH 6.5,<br>700 mM NaCl,<br>18% PEG 4000 | 15% ethylene glycol | $3PDZ1^{KSTQF}$<br>(this study) |
| $3PDZ4^{VLKSTQF}$ | 20 mM Tris-HCl, pH 8.5,<br>150 mM NaCl,<br>1 mM TCEP | 100 mM Tris pH 8.5,<br>100mM NaCl,<br>18% PEG 10000,<br>20% Glycerol | none | $3PDZ4^{KSTQF}$<br>(this study) |
| $1PDZ1^{NATRL}$ | 20 mM Tris-HCl, pH 8.0,<br>150 mM NaCl,<br>1 mM TCEP | 100mM Bicine pH 9,<br>100mM NaCl,<br>30% PEG 550 MME | none | $1PDZ1^{KSTQF}$<br>(this study) |

### Table S3. Data collection and refinement statistics for PDZ$^{KSTQF}$ structures

| | 3PDZ1$^{KSTQF}$ | 3PDZ2$^{KSTQF}$ | 3PDZ4$^{KSTQF}$ | 1PDZ1$^{KSTQF}$ |
|---|---|---|---|---|
| **PDB ID** | **9RXN** | **9RXO** | **9RXP** | **9RXQ** |
| **Beamline** | ID30A-3 (ESRF) | ID23-1 (ESRF) | ID30B (ESRF) | ID30A-3 (ESRF) |
| **Data processing** | | | | |
| Space group | P $2_12_12_1$ | P $2_12_12_1$ | P $3_221$ | P $2_12_12_1$ |
| Cell dimensions | | | | |
| *a*, *b*, *c* (Å) | 49.1, 49.9, 103.5 | 32.6, 39.1, 137.8 | 45.0, 45.0, 174.9 | 47.9, 51.5, 63.3 |
| *α*, *β*, *γ* (°) | 90.00, 90.00, 90.00 | 90.0, 90.0, 90.0 | 90.0, 90.0, 120.0 | 90.0, 90.0, 90.0 |
| Resolution range (Å) | 45 – 1.36 (1.38 – 1.36) | 35 – 1.20 (1.24 – 1.20) | 40 – 1.45 (1.54 – 1.45) | 40 – 1.38 (1.41 – 1.38) |
| Resolution limits along a*, b*, c* (Å) | N.A. | 1.17, 1.45, 1.14 | 1.45, 1.45, 1.68 | N.A. |
| Number of unique reflections | 55238 (2536) | 49076 (2304) | 30496 (1525) | 32381 (1484) |
| $R_{merge}$ | 0.101 (0.560) | 0.085 (1.415) | 0.247 (11.956) | 0.091 (0.461) |
| $R_{pim}$ | 0.016 (0.109) | 0.028 (0.479) | 0.023 (1.072) | 0.010 (0.085) |
| $I / \sigma I$ | 29.2 (5.8) | 12.1 (2.1) | 17.7 (1.5) | 50.1 (9.1) |
| $CC_{1/2}$ | 0.999 (0.949) | 0.992 (0.562) | 0.999 (0.792) | 0.999 (0.983) |
| Completeness (%) | | | | |
| spherical | 99.5 (92.8) | 81.4 (41.7) | 81.3 (24.3) | 99.3 (92.1) |
| ellipsoidal | N.A. | 94.1 (90.0) | 93.0 (50.0) | N.A. |
| Multiplicity | 39.9 (24.5) | 10.1 (9.3) | 119.1 (124.3) | 86.8 (56.3) |
| **Refinement** | | | | |
| Resolution (Å) | 45 – 1.36 | 35 – 1.20 | 39 – 1.45 | 40 – 1.38 |
| No. reflections used in refinement | 55230 | 45941 | 30492 | 32372 |
| $R_{work}$ / $R_{free}$ | 13.4 / 16.1 | 16.6 / 20.4 | 20.8 / 23.4 | 16.0 / 18.3 |
| No. of protein mol./A.U. | 2 | 2 | 2 | 2 |
| Model completeness (%) | 100 | 100 | 98.3 | 100 |
| No. atoms | | | | |
| Total Protein (hydrogens) | 3660 (1864) | 2957 (1489) | 2937 (1494) | 2917 (1479) |
| Ligands and ions | 22 (12) | 10 (6) | 0 | 7 (3) |
| Water | 441 | 274 | 128 | 244 |
| *B* factors | | | | |
| Protein | 13.32 | 22.60 | 37.00 | 20.00 |
| Ligands and ions | 25.37 | 41.95 | N.A. | 33.82 |
| Water | 27.08 | 32.09 | 39.62 | 31.00 |
| r.m.s deviations | | | | |
| Bond lengths (Å) | 0.006 | 0.009 | 0.009 | 0.010 |
| Bond angles (°) | 0.905 | 1.037 | 1.078 | 1.040 |
| Ramachandran Plot (%) | | | | |
| Favored | 98.57 | 99.44 | 98.86 | 100.00 |
| Allowed | 1.43 | 0.56 | 1.16 | 0.00 |
| Outliers | 0.00 | 0.00 | 0.00 | 0.00 |

mol./A.U.: molecule per assymetric unit

**Table S4. Data collection and refinement statistics for PDZ$^{VLKSTQF}$ and 1PDZ1$^{NATRL}$ structures**

| | **3PDZ1$^{VLKSTQF}$** | **3PDZ4$^{VLKSTQF}$** | **1PDZ1$^{NATRL}$** |
|---|---|---|---|
| **PDB ID** | **9RXR** | **9RXS** | **9RXT** |
| **Beamline** | BM07 | BM07 | ID30A-3 |
| **Data processing** | | | |
| Space group | $P3_1$ | $P4_32_12$ | $P2_13$ |
| Cell dimensions | | | |
| *a*, *b*, *c* (Å) | 36.0, 36.0, 72.0 | 52.4, 52.4, 195.8 | 64.2, 64.2, 64.2 |
| *α*, *β*, *γ* (°) | 90.0, 90.0, 120.0 | 90.0, 90.0, 90.0 | 90.0, 90.0, 90.0 |
| Resolution range (Å) | 35 – 2.11 (2.15 – 2.11) | 49 – 2.00 (2.13 – 2.00) | 40 – 1.36 (1.38 – 1.36) |
| Resolution limits along a*, b*, c* (Å) | N.A. | 2.47, 2.47, 1.79 | N.A. |
| Number of unique reflections | 5710 (285) | 12585 (629) | 19221 (904) |
| $R_{merge}$ | 0.316 (1.843) | 0.150 (2.320) | 0.125 (3.780) |
| $R_{pim}$ | 0.103 (0.611) | 0.021 (0.520) | 0.011 (0.417) |
| $I / \sigma I$ | 6.2 (2.0) | 20.8 (2.7) | 30.3 (1.5) |
| $CC_{1/2}$ | 0.979 (0.486) | 1.000 (0.878) | 1.00 (0.584) |
| Completeness (%) | | | |
| spherical | 94.7 (78.3) | 65.0 (19.4) | 99.8 (97.7) |
| ellipsoidal | N.A. | 90.2 (86.4) | N.A. |
| Multiplicity | 10.3 (9.7) | 48.9 (41.7) | 120.0 (75.7) |
| **Refinement** | | | |
| Resolution (Å) | 24 – 2.11 | 25 – 2.00 | 37 – 1.36 |
| No. reflections used in refinement | 5690 | 12547 | 19171 |
| $R_{work}$ / $R_{free}$ | 17.3 / 22.8 | 19.8 / 24.2q | 15.7 / 19.7 |
| No. of protein mol./A.U. | 1 | 2 | 1 |
| Model completeness (%) | 100 | 99.5 | 100 |
| No. of atoms | | | |
| Protein all (hydrogen) | 1786 (902) | 2889 (1464) | 1519 (770) |
| Ligands and ions all (hydrogen) | 0 | 14 (8) | 11 (16) |
| Water | 106 | 115 | 148 |
| *B* factors | | | |
| Protein | 18.26 | 47.17 | 25.02 |
| Ligands and ions | N.A. | 58.18 | 26.02 |
| Water | 19.59 | 39.51 | 33.88 |
| r.m.s deviations | | | |
| Bond lengths (Å) | 0.007 | 0.010 | 0.010 |
| Bond angles (°) | 0.651 | 0.744 | 0.994 |
| Ramachandran Plot (%) | | | |
| Favored | 98.13 | 98.88 | 98.86 |
| Allowed | 1.87 | 1.12 | 1.14 |
| Outliers | 0.00 | 0.00 | 0.00 |
